# Neonate personality affects early-life resource acquisition in a large social mammal

**DOI:** 10.1101/2022.03.15.484465

**Authors:** Bawan Amin, Dómhnall J. Jennings, Alison Norman, Andrew Ryan, Vasiliki Ioannidis, Alice Magee, Hayley-Anne Haughey, Amy Haigh, Simone Ciuti

**Author notes:** **Author Contributions**, BA, DJJ and SC conceived the ideas and designed methodology; BA, AN, AR, VI, AM, HAH and AH collected the data; AN, AM and HAH carried out the data scoring; BA and SC analysed the data; BA wrote the manuscript, revised by DJJ and SC. All authors contributed critically to the drafts and gave final approval for publication.

## Abstract

Current debate in the field of animal personality revolves around whether personality is reflecting individual differences in resource allocation or acquisition. Despite the large body of literature, the proximate relationships between personality, resource allocation, and acquisition are still unclear, especially during early stages of development. Here we studied how among-individual differences in behaviour develop over the first 6 months of life, and their potential association with resource acquisition in a free-ranging population of fallow deer (*Dama dama*). We related proxies of neonate personality – i.e. neonate physiological (heart rate) and behavioural (latency to leave at release) anti-predator responses to human handling – to the proportion of time fawns allocated to scanning during their first summer and autumn of life. We then investigated whether there was a trade-off between scanning time and foraging time in these juveniles, and how it developed over their first 6 months of life. We found that neonates with longer latencies at capture (i.e. risk-takers) allocated less time scanning their environment, but that this relationship was only present when fawns were 3-6 months old during autumn, but not when fawns were only 1-2 months old during summer. We also found that time spent scanning was negatively related to time spent foraging – a relationship rarely tested in juveniles of large mammals - and that this relationship becomes stronger over time, as fawns gradually switch from a nutrition rich (milk) to a nutrition poor (grass) diet. Our results highlight a potential mechanistic pathway in which neonate personality may drive differences in early-life resource acquisition, through allocation, of a large social mammal.

## Introduction

Individuals tend to differ in their average behaviour and these among-individual differences, when consistent over time and across context (i.e. “animal personality”), have been shown to play an important role in ecology and evolution (Wolf & Weissing, 2012). Current theory such as the extended pace-of-life syndrome hypothesis (POLS) suggests that among-individual differences in behaviour mediate within-species differences in life-history strategies on a fast-slow continuum (Réale et al., 2010). Within this framework, behaviour, physiology and life-history are expected to covary. Individuals with a faster pace-of-life (POL) are expected to be bolder, more active, and to allocate more resources to growth and short-term reproduction than individuals with a slower POL (Réale et al., 2010). This increased resource allocation in current growth or reproduction is predicted to come at the cost of survival: animals with a faster POL are expected to have a shorter lifespan relative to those with a slower POL (Réale et al., 2010).

Over the last decade, empirical research has not provided conclusive results for the main predictions of POLS (Laskowski et al., 2021; Moiron et al., 2020; Royauté et al., 2018), suggesting that there is a greater complexity than expected in the covariation between behaviour and life-history. Recently, Laskowski et al. (2021) suggested that such a covariance may vary depending on the interaction between among-individual variation in behaviour and actual resource acquisition. If among-individual variation in behaviour is closely related to resource acquisition, with bolder animals gaining even more resources than shyer ones, then the trade-off between survival, current growth and reproduction may be weakened or even disappear (Laskowski et al., 2021). This is also what recent meta-analyses seem to suggest (Moiron et al., 2020; Haave-Audet et al., 2021). Neither of these meta-analyses found support of resource allocation as main driver of individual variation, suggesting that resource acquisition may instead be the main driver of among-individual variation (Haave-Audet et al., 2021).

Additional complexity, often leading to contrasting evidence in animal personality science, is associated with changes of behaviour over different life-stages (Stamps & Groothuis, 2010; Cabrera et al., 2021). Although there is clear evidence that individuals behave in a consistent way within life-stages, including the earliest stages of life (Amin et al., 2016; Fucikova et al., 2009; Guenther & Trillmich, 2015; Dhellemmes et al., 2020), the same cannot be said of individual consistency across different life-stages (see Cabrera et al., 2021 for a review and references therein). Furthermore, the limited number of studies that have investigated among-individual differences across life-stages have either done so on captive populations (Wuerz & Krüger, 2015; Neave et al., 2020; Favati et al., 2016), or have measured behaviour only during capture (Class & Brommer, 2015; Petelle et al., 2013) or within artificial settings (Kelley et al., 2015; Hall et al., 2015). There is a paucity of studies that have investigated whether these traits measured in controlled settings are actually related to life history in the wild (Niemelä & Dingemanse, 2014).

Consequently, the relationship between animal personality and life history related traits during early stages of maturation and development in a wild setting has yet to be tested. To address this shortcoming, we tested whether repeatable among-individual differences were associated with behavioural strategies affecting early-life resource acquisition in a free-living population of fallow deer fawns (*Dama dama*). These were monitored from birth to 6 months, through the key transition from solitary to group-living life. Fallow deer are a hider species (Lent, 1974): fawns experience a solitary life during their first 2-4 weeks of life remaining hidden in vegetation while occasionally being visited by their mother for maternal care (Chapman & Chapman 1997). We recently showed that repeatable among-individual differences are present during the first days of life in this population (Amin et al., 2021). Some neonates display repeatable active responses – i.e. elevated heart rates and short latency to leave when released - whereas other neonates are bolder and less risk aversive – i.e. they maintain low heart rates during human handling and have longer latencies to leave once released (Amin et al., 2021).

A few weeks after birth, most fawns make the transition from a solitary life to a group-living one and they join the female herd with their mothers, gradually shifting from a milk-based diet to a fully independent herbivorous diet (Chapman & Chapman 1997). When they join the main herd with their mothers, fawns are expected to trade-off their time budgets, as typical for herbivores, between anti-predator behaviour, i.e. scanning the landscape for potential threats, and resource acquisition, i.e. foraging (Sih, 1980; Lima, 1987; Bachmann, 1993). Simulations have recently shown that scanning dictates the amount of resources acquired and not vice versa (Sirot et al., 2021). Since bolder individuals are predicted to spend less time scanning (Bergvall et al., 2011; Uchida et al., 2019), scanning behaviour could act as a feature of personality that in turn dictates resource acquisition. By allocating less time in scanning, individuals may be able to increase resource acquisition. Shedding light on these relationships will therefore provide a mechanism in which personality, through allocation, explains resource acquisition during the early stages of independence in juveniles.

Here we tested whether neonate personality of fallow deer fawns, recorded during their hider phase, is related to the time they allocate to scanning while living in a group, during their first 6 months of life. In order to do that, we first tested whether scanning times were repeatable between individuals. Our main hypothesis then was that among-individual differences in neonate traits would be covary with among-individual differences in scanning time. Specifically, we predicted that animals who react more boldly at capture, i.e. lower heart rates and longer latencies, also behave more boldly while in the herd, i.e. spend less time scanning. We then tested whether time spent scanning is inversely related to foraging time, our proxy for resource acquisition. Although this relationship is fairly clear in adults across vertebrates (Caro, 2005), juveniles have been shown to scan the environments less than adults in several bird and mammal species (see Caro, 2005 for a review), and could therefore also differ in their time budget trade-offs. We predicted that the trade-off between scanning and foraging would be present in fawns, and furthermore, that it would increase in strength when fawns grow older as they switch from a maternally provisioned nutrition rich (i.e. milk) to a nutritionally poorer (i.e. grazer) diet (Arenz & Leger, 2000).

## Methods

### Study site and study population

This study was conducted on a population of European fallow deer resident in the Phoenix Park, a 7 km^2^ enclosed park located near the centre of Dublin, Ireland (53.3559° N, 6.3298° W). Vegetation in the park is predominately open grassland (~80%) with the remaining area composed by mixed woodland. Our study population of deer was estimated to be over 600 individuals over the course of this study (late summer estimates after the fawning). The majority of fawns are born from early June to early July. Fallow deer are a hider species and fawns remain hidden, usually in tall grass or understory vegetation, away from the main doe herd during the first two-three weeks of life following which they are brought into the doe herd by their mothers (Chapman & Chapman, 1997; Ciuti et al., 2006). The only natural predator present in the park is the red fox (*Vulpes vulpes*), although fawns are also occasionally preyed upon by unleashed domestic dogs (*Canis lupus familiaris*). Deer are culled annually by professional stalkers over the winter period as part of the population management led by the Office of Public Works.

### Neonate captures

Fawns have routinely been captured and ear-tagged with unique numbered and coloured plastic tags (Allflex medium, Mullinahone Co-op, Ireland) since the early 1970’s as part of the monitoring and management of the herd (Hayden et al. 1992). Fawns were located by patrolling geographical areas traditionally used by does as fawning sites daily in June, when the majority of the births happen. Using fishing nets (1-1.5m diameter; various brands), we located and tagged a total of 185 fawns over two consecutive years (n = 102 in 2018, n = 83 in 2019), of which 91 were recaptured once (n = 43 in 2018, n = 48 in 2019), 33 twice (n = 14 in 2018, n = 19 in 2019), and 9 three times or more (n = 4 in 2018, n = 5 in 2019). We recorded the following confounding variables which have been shown to affect neonatal response to handling (Amin et al. 2021): weight (in kg) was measured using a digital scale by laying the fawn in a 100-litre bag (resolution: 0.01 kg – Dario Markenartikelvertrieb, Hamburg, Germany); air temperature was measured at the bed-site location using a digital thermometer (Grandbeing, China). We quantified the behaviour of the fawn prior to capture (prior behaviour) by recording whether the fawn was in motion (yes = 1, no = 0), turned its head to look around (yes = 1, no = 0), kept its head up or down (up = 1, down = 0), had its ears up or down (up = 1, down = 0), was down but got up (yes = 1, no = 0), and attempted to run away (yes = 1, no = 0). We took the mean of all these scores as a measure of prior behaviour, where 1 indicated the most active behaviour and 0 the least active behaviour (*sensu* Amin et al., 2021).

Directly relevant to this study, we selected a physiological trait (heart rates prior to release, i.e. a physiological response of fawns to human handling) and a behavioural trait (latency to leave upon release), both shown to be repeatable at the among-individual level previously (Amin et al., 2021). Heart rates were taken directly before the weighting of the fawns and quantified by counting the number of beats per 20 seconds using a Lightweight Dual Head Stethoscope (MDF®, California, USA). The latency to leave (in seconds) on release was defined as the time it took the fawn to stand up after opening the weighing bag. We took 10 seconds as the maximum value and assigned that to individuals that had not moved before then (Amin et al., 2021).

### Focal observation in the herd

Time budgets were computed from focal sampling during summer and autumn in each year. Summer data collection took place in July and August of each year when newborn fawns join the female herd for the first time. Although the timing of emergence into the herd can be variable between individuals, most fawns make their first appearances in the herd in the summer months (See Supplementary S1). Autumn data collection took place from mid-September until early December, overlapping with the rutting season. The temporal overview of the different data collection periods is displayed in Fig.1.

**Fig 1:**
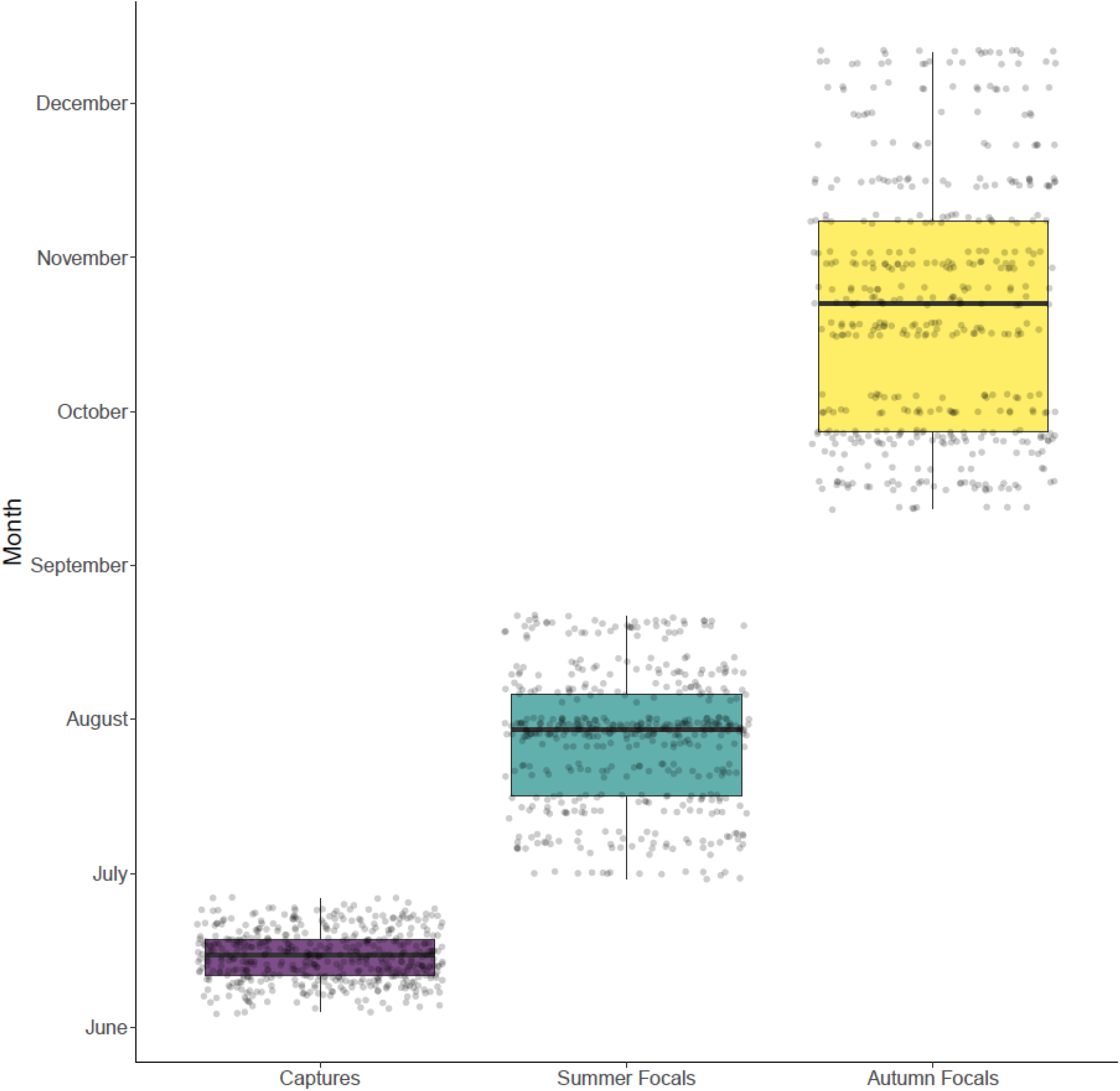
A temporal overview of the different data collection periods in 2018 and 2019, which were the neonate captures and the focal observations taken in summer and autumn. Jittered points indicate individual observations.

Observations were taken between 09.00-17.00 hours, generally in dry weather with high visibility. Focal subjects were observed using a spotting scope, at a distance no closer than 50 m, allowing the observer to maintain their distance and minimize their impact on the fawns’ behaviour. Sampling of the focal individuals was random, with a priori determined rotation system used to find and sample most fawns available in the herd. We walked through the female day range and identified and selected a social group with active, non-resting fawns. A group was defined as multiple clustered individuals that were within 50 m of each other. If a group included multiple active fawns, we selected the focal individual randomly. At the start of the observation, we recorded the total number of deer in that group. Groups were loosely aggregated, with constant fission-fusion throughout the day. The fawn was continuously observed for up to 25 minutes. Focals were often ended early due to fawns moving out of sight, i.e. laying down in the long grass, entering a traveling bout with the group, or a major herd disturbance. The fawn’s behaviour during this period was recorded on a Dictaphone (Olympus VN-540PC) and transcribed later. The mean observation duration, excluding time out of sight, was 3.74 (SD = 3.97) minutes for the summer and 7.66 (SD = 5.81) minutes for the autumn season.

For each focal observation, we recorded the fawn’s position in the group at the start and end, based on the number of deer between the fawn and the edge of the group. This ranged from 0 to 3, with 0 being the fawn at the outermost edge of the herd, and 3 being three or more deer between the fawn and edge of the herd. We did so in order to account for potentially increased scanning rates of individuals observed at the edge of a group (Caro, 2005). As a measure of human disturbance – and its potential effect on scanning rates (Ciuti et al., 2012), we also recorded the total amount of park visitors within a 50 m radius of the focal fawn during the observation. Once all active fawns in a herd had been recorded, the observers moved on and another herd was selected. We avoided resampling the same fawns during the same session, unless the first observation was very short (< 2 minutes). Initially, fawns were chosen at random. As more focals were obtained, fawns were chosen more selectively, prioritizing individuals with a lesser number of observations. In total, we collected 477 focal observations on 137 fawns during summer and 430 focal observations on 145 fawns during autumn, making up a total of 907 focal observations with a total cumulative duration of 84.6 hours.

Time budgets were extracted from the audio recordings using Jwatcher (Version 1.0) software. Twenty-six behaviours were recorded, a full description of each behaviour can be found in Table S1. Proportion of time spent on each behaviour was calculated from the total time of each observation, excluding time spent out of sight. Proportion of time spent scanning, defined as standing still with the head above the shoulder height, was used as a measure of scanning time. We accumulated the time spent scanning while chewing and without chewing, since it was difficult to distinguish the two behaviours in the field. Proportion of time spent foraging was calculated by combining the proportions of time spent grazing, defined as unselectively feeding on grass and ground vegetation with the head below the shoulders, and browsing, defined as selectively feeding on leaves, bark and top of plants (see Supplementary ST1 for full definitions).

### Ethical note

Captures and handling were carried out giving the highest priority to animal welfare. Fawns that were evidently newborn (a fully wet coat) were not captured and in such instances, we abandoned searches in that area to avoid disturbing the fawn. Gloves were always worn during handling to prevent transfer of human odours to the fawn (Galli et al, 2008). We operated in silence during animal handling and left the bed-site immediately after the release of the fawn. Fawns were released in a location adjacent to the capture site and facing in a direction away from the capture team. The capture, handling, tagging and sampling of fawns was supervised by a certified and experienced wildlife biologist. Regular monitoring of the tagging regime has shown there are no survival implications in this population (see also Hayden et al., 1992). The focal data collection was observational: observers kept a minimum distance of 50m from the deer to avoid disturbing their behaviour. The study protocol and all research procedures were approved by the Animal Research Ethics Committee (University College Dublin) under permit number AREC-E-18-28. All methods were in accordance with the Guidelines for the treatment of animals in behavioural research and teaching (Animal Behaviour, 2020).

### Statistical Analysis

All analyses were conducted using R 3.5.1 (R Core Team, 2020). To give a general overview of the analyses expanded upon below, repeatability of scanning and the covariance between neonate traits and time spent scanning were examined using bivariate mixed effect models (Houslay & Wilson, 2017). We then analysed the trade-off between scanning and foraging using univariate mixed-effect models. All response variables and numerical explanatory variables were scaled prior to analysis, such that each variable was centred at their mean value and standardised to units of 1 phenotypic standard deviation. This has been recommended to improve model convergence and result interpretation (Houslay & Wilson, 2017). Full details of the statistical analysis are provided below in the subsections.

### Neonate traits at capture and scanning time

To estimate the repeatability of and the among-individual covariation between the neonate traits at capture (heart rate and latency to leave) and scanning time while in the herd (in summer and autumn separately), we used multivariate mixed models, under a Bayesian MCMC framework, which are regarded as the state-of-the-art method for personality research (Houslay & Wilson, 2017; Dingemanse & Wright, 2020; Hertel et al., 2020). Multivariate mixed models were fitted via the *MCMCglmm*-package (Hadfield, 2010). To determine whether either heart rate or latency to leave at capture were correlated with scanning behaviour, we fitted four separate bivariate mixed models. Two of the models had heart rate and scanning time as response variables, one model for the summer period and the other for the autumn period. The other two models had latency to leave and scanning time as response variables, with also one model for the summer period and one model for the autumn period. Within-individual covariation between the two responses of each bivariate model were set to 0, since the two responses within each model were not measured at the same time (see Hadfield, 2010). Correlation coefficients at the among-individual level (r_i_) and repeatability estimates, along with their 95% credible intervals, were computed following Houslay & Wilson (2017). In all models, Fawn ID was included as random intercept. For each bivariate model, we included only individuals that had at least one datapoint per response variable. We also omitted rows with missing values in any of the explanatory variables from the analysis.

In all cases we used a weakly informative prior (*R = list(V = diag(2), nu = 0.002; G = list(G1 = list(V = diag(2), nu = 1.002))*). The neonate response variables (heart rate and latency to leave) were log-transformed prior to analysis to improve model fit and meet model assumptions regarding the gaussian distribution of errors. The scanning time response variable was in all cases logit-transformed, i.e. log(y/[1 - y]), as suggested by Warton & Hui (2011). Since the logit of 0 and 1 translate to −∞ and ∞, we added the smallest non-zero value to both the numerator and denominator of the logit equation (Warton & Hui, 2011). We used *a priori* model structures for each response variable, which in the case of the neonate capture traits were based on a previous study (Amin et al., 2021). In the case of scanning time, we included explanatory variables that contained information on the context of each observation, where we included both the linear and quadratic terms for all the numerical explanatory variables to allow non-linear effects. To avoid overfitting of the model, we simplified the full model by only removing the quadratic term of a variable when pMCMC>0.1. The final model structures for each model are given in Table 1 and Table 2, where the columns indicate the response variables and the rows the explanatory variables.

**Table 1:** Structure and output of the final bivariate models (MCMCglmm) used for the analysis of the among-individual covariation between heart rate at capture and time spent scanning in the summer (*Heart rate-scanning summer model*) as well the covariation between latency to leave at capture and time spent scanning in the summer (*Latency-scanning summer model*). Posterior means with their associated 95% Credible Intervals of each of the explanatory variables (rows) included are given. Empty cells indicate that the explanatory variable was not included in the model for the respective response variables (model structures for the neonate traits defined by Amin et al. 2021). The position in the herd was not taken during the summer of 2018 and therefore, left out of the two models that were used for the summer season.

| Variable | Posterior mean [95% CrI] |  |  |  |
| --- | --- | --- | --- | --- |
|  | Heart rate-scanning summer model |  | Latency-scanning summer model |  |
|  | Heart rate | Scanning (summer) | Latency | Scanning (summer) |
| Intercept | -0.164 [-0.356, 0.061] | 0.033 [-0.153, 0.216] | -0.649 [-1.305, -0.093] | 0.030 [-0.142, 0.209] |
| Prior behaviour | 0.087 [0.007, 0.163] |  | -0.127 [-0.198, -0.051] |  |
| Prior behaviour<sup>2</sup> | -0.069 [-0.147, 0.012] |  |  |  |
| Weight | 0.220 [0.144, 0.301] |  | -0.142 [-0.218, -0.056] |  |
| Weight<sup>2</sup> | 0.033 [-0.039, 0.109] |  | 0.064 [-0.004, 0.136] |  |
| Year (2019) |  |  | 0.099[-0.154, 0.372] |  |
| Capture |  |  | -0.172 [-0.330, -0.031] |  |
| Time of day | 0.127 [-0.002, 0.245] | -0.159 [-0.252, -0.060] |  | -0.164 [-0.264, -0.072] |
| Time of day<sup>2</sup> | -0.000 [-0.102, 0.105] |  |  |  |
| Air temperature | 0.080 [0.004, 0.154] |  |  |  |
| Sex (m) | 0.238 [-0.011, 0.501] | -0.319 [-0.555, -0.115] |  | -0.308 [-0.522, -0.087] |
| Season (2018) |  | -0.421 [-0.653, -0.176] |  | -0.418 [-0.658, -0.191] |
| Number of people |  | 0.072 [-0.001, 0.150] |  | 0.073 [-0.005, 0.151] |
| Group size |  | 0.088 [0.003, 0.163] |  | 0.088 [0.004, 0.159] |
| Birthday (in days) |  | 0.031 [-0.054, 0.118] |  | 0.027 [-0.057, 0.115] |
| Days since emergence |  | -0.267 [-0.399, -0.125] |  | -0.265 [-0.404, -0.135] |
| Observation length (ms) |  | 0.159 [0.078, 0.247] |  | 0.160 [0.071, 0.241] |
| Observation length (ms)<sup>2</sup> |  | -0.173 [-0.256, -0.092] |  | -0.175 [-0.249, -0.093] |

**Table 2:** Structure and output of the final bivariate models (MCMCglmm) used for the analysis of the among-individual covariation between heart rate at capture and time spent scanning in the autumn (*Heart rate-scanning autumn model*) as well the covariation between latency to leave at capture and time spent scanning in the autumn (*Latency-scanning autumn model*). Posterior means with their associated 95% Credible Intervals of each of the explanatory variables (rows) included are given. Empty cells indicate that the explanatory variable was not included in the model for the respective response variables (model structure for neonate traits defined by Amin et al. 2021).

| Variable | Posterior mean [95% CrI] |  |  |  |
| --- | --- | --- | --- | --- |
|  | Heart rate-scanning autumn model |  | Latency-scanning autumn model |  |
|  | Heart rate | Scanning (autumn) | Latency | Scanning (autumn) |
| Intercept | -0.145 [-0.342, 0.049] | -0.036 [-0.253, 0.186] | -0.969 [-1.554, -0.450] | -0.033 [-0.245, 0.193] |
| Prior behaviour | 0.093 [0.021, 0.179] |  | -0.135 [-0.214, -0.060] |  |
| Prior behaviour<sup>2</sup> | -0.041 [-0.128, 0.040] |  |  |  |
| Weight | 0.188 [0.112, 0.273] |  | -0.087 [-0.178, -0.004] |  |
| Weight<sup>2</sup> | 0.034 [-0.049, 0.106] |  | 0.066 [-0.008, 0.142] |  |
| Year (2019) |  |  | 0.139 [-0.127, 0.399] |  |
| Capture |  |  | -0.249 [-0.391, -0.107] |  |
| Time of day | 0.117 [-0.000, 0.239] | -0.148 [-0.252, -0.043] |  | -0.146 [-0.253, -0.044] |
| Time of day<sup>2</sup> | 0.016 [-0.115, 0.129] |  |  |  |
| Air temperature | 0.079 [-0.001, 0.155] |  |  |  |
| Sex (m) | 0.221 [-0.024, 0.485] | -0.077 [-0.306, 0.154] |  | -0.083 [-0.313, 0.116] |
| Season (2018) |  | 0.304 [0.063, 0.566] |  | 0.295 [0.056, 0.550] |
| Number of people |  | 0.048 [-0.031, 0.119] |  | 0.050 [-0.024, 0.123] |
| Group size |  | 0.050 [-0.033, 0.129] |  | 0.054 [-0.027, 0.130] |
| Group size<sup>2</sup> |  | 0.075 [-0.003, 0.159] |  | 0.074 [-0.010, 0.153] |
| Birthday (in days) |  | -0.004 [-0.097, 0.081] |  | -0.010 [-0.097, 0.077] |
| Position in the herd |  | 0.039 [-0.037, 0.121] |  | 0.035 [-0.041, 0.113] |
| Position in the herd<sup>2</sup> |  | 0.089 [0.003, 0.169] |  | 0.088 [0.005, 0.172] |
| Days since emergence |  | -0.238 [-0.369, -0.104] |  | -0.238 [-0.366, -0.115] |
| Observation length (ms) |  | 0.013 [-0.081, 0.110] |  | 0.022 [-0.079, 0.118] |

All MCMC-chains were run for a total length of 1,050,000 iterations, with a thinning of 500 and a burnin of the first 50,000 iterations, leading to a total of 2,000 saved iterations. Model convergence was checked by running 4 separate chains for each bivariate model and calculating the multivariate scale reduction factor (Brooks & Gelman, 1998), which never exceeded 1.1. We also visually inspected the chains, ensuring that every parameter had an effective sample size of at least 1,000, and the autocorrelation of the posterior means and variances. From these, we concluded that the chains had converged properly and had negligible autocorrelations. Inferences concerning each of the correlations were made based on the posterior mean and the highest posterior density interval. We considered a relationship to be meaningful if less than 5% of the posterior distribution crossed zero (Allen et al., 2017; Jennings et al., 2018). To visualise the relationships between the responses of the bivariate, we extracted the posterior means of the random intercepts (BLUPs; Houslay & Wilson 2017). Full details on the bivariate models, including all the code, model summaries and model diagnostics are given as supplementary material (Supplementary S2).

### Trade-off between scanning and foraging

To investigate the possible trade-off between scanning and foraging in young fawns and the possible change over ontogeny, we fitted a linear mixed-effect model (*lme4* package, Bates et al. 2015). Time spent scanning and time spent foraging were quantified as proportions of total time, which were then logit-transformed (Warton & Hui, 2011). Since scanning is proposed to be driving resource acquisition (Sirot et al., 2021), we used foraging time as our response variable and scanning time as explanatory variable. To investigate change over time, we included the day of the year as a numerical explanatory variable, along with its interaction with scanning time. We included the quadratic terms of scanning time and day of the year to allow for non-linear effects. Finally, to correct for the effect of observation length on our estimates of foraging behaviour (see sensitivity analysis below; supplementary S3), we also included the duration of each observation as an explanatory variable. The predicted model effect following from this model was visualized using the *effects-package* with 95% marginal confidence intervals (Fox & Weisberg, 2018; 2019).

### Sensitivity analysis

Initially, we aimed to include foraging time as a response variable in our bivariate models as well, in addition to scanning time and the neonate response variables, to investigate the relationship between neonate personality and resource acquisition directly. Prior to running our bivariate models, however, we investigated the stability of foraging time and scanning time estimates over different observation lengths. This was done because very short observations may produce biased time budgets (Childress and Lung, 2003). For that purpose, we ran a sensitivity analysis (Supplementary S3). From the sensitivity analysis we concluded that foraging time was strongly affected by observation length and failed to stabilise even with increasing observation lengths. We therefore decided not to include foraging time as a response variable in our bivariate models, which we use to estimate among-individual covariation. Scanning time, on the other hand, was relatively robust and, especially in autumn, barely affected by observation length. There was some minor underestimation of scanning time for very short observations, mainly during summer, and we therefore included observation length as an explanatory variable in our bivariate models for scanning time.

## Results

### Neonate traits at capture and scanning time

Both neonate traits measured at capture were found to be repeatable among individuals (heart rate: R = 0.35, 95% CrI [0.18, 0.51], N = 145 individual fawns; latency to leave: R = 0.33, 95% CrI [0.17, 0.48], N = 145). The proportion of time that fawns spent scanning was also repeatable among individuals, in both summer as well as autumn (summer: R = 0.12, 95% CrI [0.06, 0.18], N = 137; autumn: R = 0.17, 95% CrI [0.09, 0.25], N = 145). The posterior means and 95% CI of the explanatory variables used for estimating repeatability and among-individual covariance between neonate traits at capture and time spent scanning are given in Table 1 (summer models) and Table 2 (autumn models). We found no meaningful relationship between heart rates and scanning time in summer nor in autumn (Table 3; Fig. 2A; 2C). There was also no clear pattern between latency to leave and scanning time in the summer (Table 3; Fig. 2B). In autumn, however, we did find a meaningful negative relationship between latency to leave and scanning time. Individuals with higher latencies to leave as neonates in June spent less time scanning their environment in autumn (Table 3; Fig. 2D).

**Table 3:** Correlations between different traits, at the among-individual level, extracted from bivariate models. Correlations displayed in bold indicate statistically meaningful effects.

| Response 1 | Response 2 | Correlation coefficient ($r_i$) | 95% Credible intervals | $N_{\text{fawns}}$ |
| --- | --- | --- | --- | --- |
| Heart rate | Scanning (summer) | -0.169 | [-0.526, 0.207] | 137 |
| Latency | Scanning (summer) | -0.024 | [-0.434, 0.354] | 137 |
| Heart rate | Scanning (autumn) | 0.014 | [-0.355, 0.417] | 145 |
| Latency | Scanning (autumn) | **-0.359** | **[-0.669, -0.028]** | 145 |

**Fig 2:**
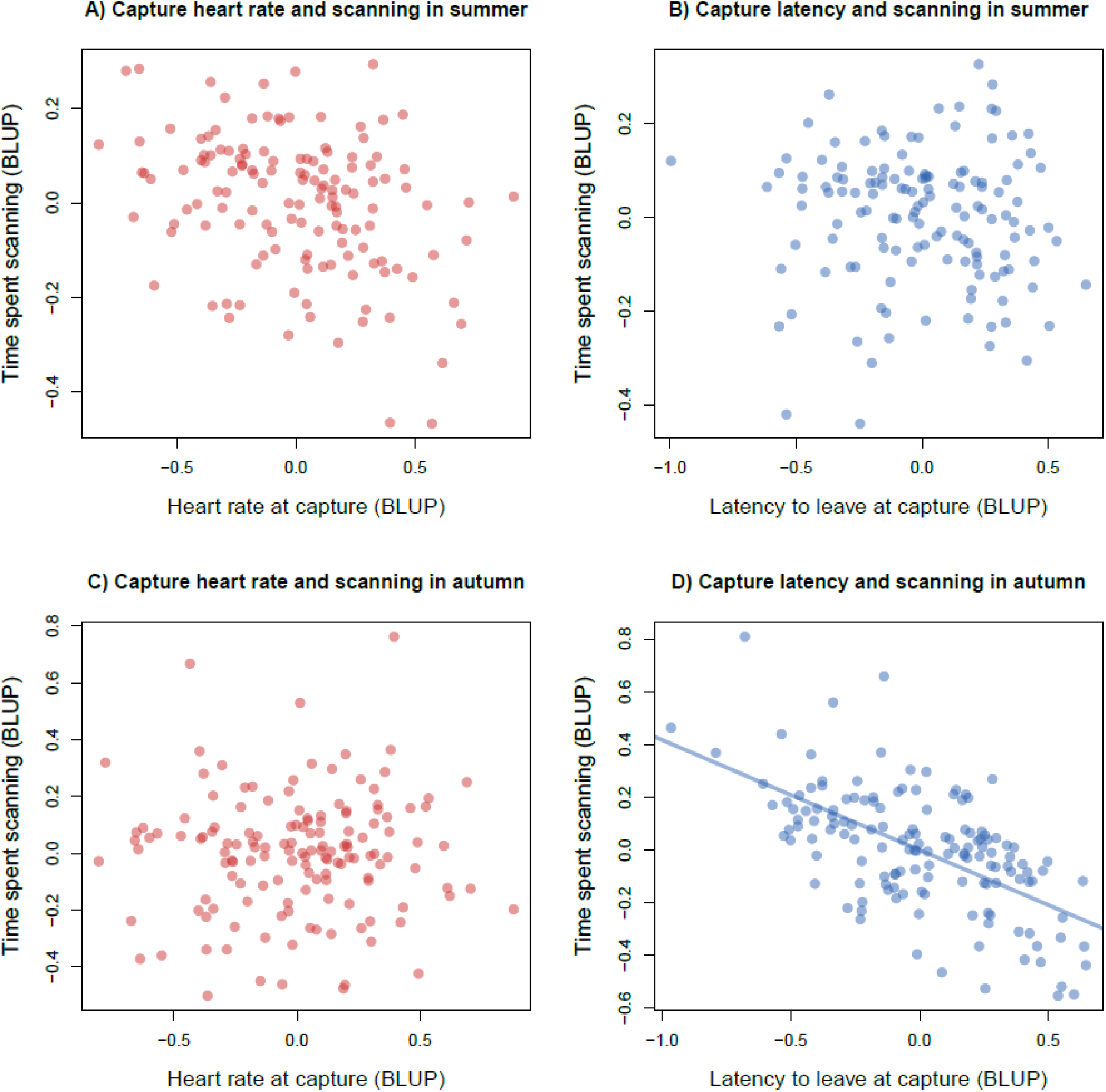
The relationships between A) heart rate and scanning time in summer, B) latency to leave and scanning time in summer, C) heart rate and scanning time in autumn and D) latency to leave and scanning time in autumn. Posterior means of the random intercepts (BLUPs) were used here for visualisation purposes only. Solid trendline indicates a meaningful effect.

### Trade-off between scanning and foraging

Fawns decreased their scanning time and increased their foraging time as they aged (see Fig. 3), i.e. during the switch from a milk-based to a grazer diet (fully weaned). Suckling events per hour (s/h) were indeed high in summer (focal observations: 0.81 suckling/hour, range per month: 0.57-0.94 s/h) and nearly disappeared in autumn (focal observations: 0.15 s/h, range per month: 0.00-0.25 s/h). We investigated whether there was a trade-off between time spent scanning and time spent foraging and whether and how this developed over time. Time spent scanning negatively affected time spent foraging (linear term: β = −0.90 ± 0.06 SE, p < 0.001, N = 907 focal observations on N = 156 fawns; quadratic term: β = −0.47 ± 0.06 SE, p < 0.001, N = 907 focal observations on N = 156 fawns) and this association only became stronger over time (Fig. 4), given the strong negative effect of the interaction between scanning time and days of the year (β = −0.20± 0.02 SE, p < 0.001, N = 907 focal observations on N = 156 fawns).

**Fig. 3.**
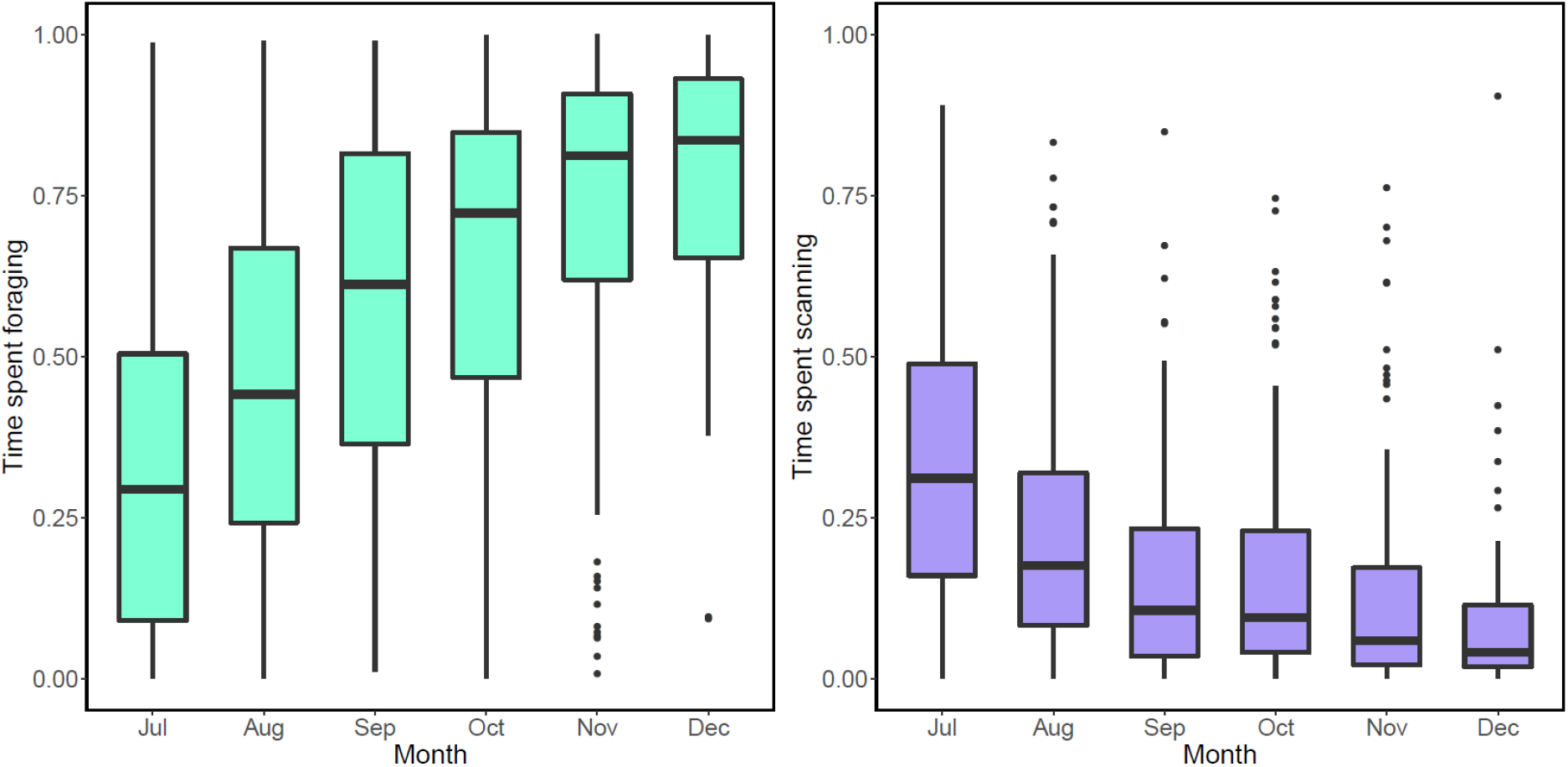
The increase in time spent foraging (left plot) and decrease in time spent scanning (right plot) of fallow deer fawns over the first 6 months of life. The times spent are given as proportions of total time of active bouts while in a group of deer.

**Fig. 4:**
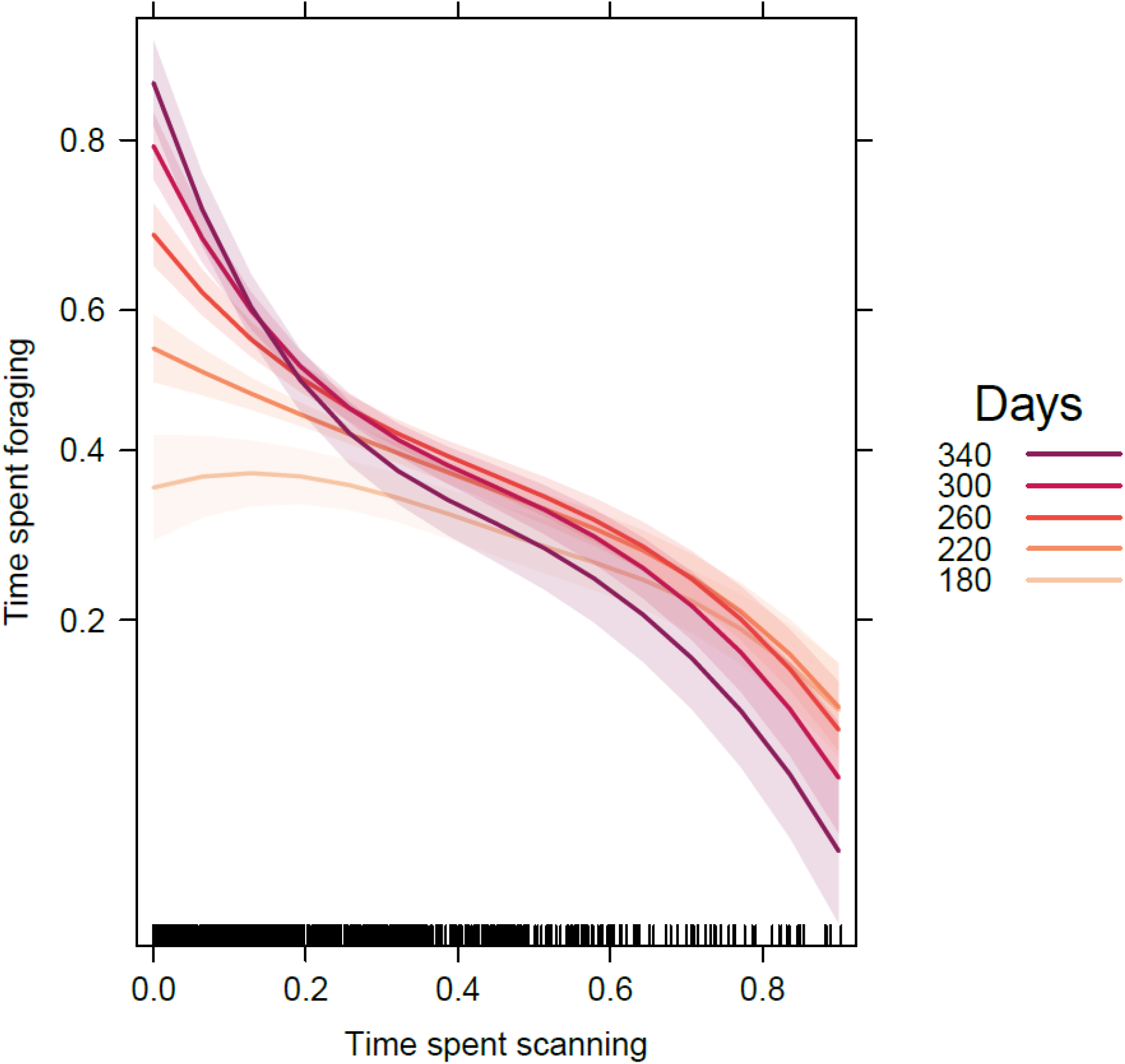
The relationship between the proportion of time spent scanning and the proportion of time spent foraging over time, i.e., when fawns gradually moved from a mainly milk-based diet to a grazer one (fully weaned). Predicted patterns (lines) are surrounded by marginal 95% confidence intervals (shaded polygons). Different time periods are indicated by different colours, with dates later in the year being represented by darker colours (day 180 = 29 June; day 220 = 8 August; day 260 = 17 September; day 300 =27 October; day 340 = 6 December)

## Discussion

Current debate within the field of animal personality focuses on whether among-individual differences in behaviour reflect life-history differences in allocation or acquisition (Laskowski et al., 2021). The theoretical framework, however, does not address the ontogeny of animal personality, which may be why the empirical support for predictions of main theories such as the extended POLS has been ambivalent (Royauté et al., 2018; Moiron et al., 2020). The ontogeny of among-individual difference remains understudied, especially in wild populations (Bell et al., 2009; Cabrera et al., 2021). Here we provide novel insights on the development of among-individual differences in juveniles of a wild large mammal, by studying fallow deer fawns from birth until their sixth month of life. In line with our predictions, we found that repeatable among-individual differences in behavioural response of neonates were related to the time they allocated to scanning their environments while in the herd with their conspecifics, which also was repeatable among individuals. This scanning behaviour was negatively related to time spent foraging, and this relationship only got stronger over time, suggesting among-individual differences in resource acquisition, through among-individual differences in time allocation. Contrary to our expectations, however, the relationship between neonate traits and time spent scanning was only present in autumn, but not earlier in summer, and also only involved the behavioural neonate response (i.e. latency to leave), and not the physiological response (i.e. heart rate). Altogether, our results show that among-individual differences are present shortly after birth and that these differences likely drive resource acquisition months later. This highlights a potential mechanistic pathway in which among-individual differences may lead to differences in resource acquisition in the earliest stages of maturation. These correlates between behaviours can, however, weaken or even diminish during major transitional phases in life-history in the wild. These results provide novel insights into the theory on animal personality, by showing that resource allocation, in this case through time budgets, and resource acquisition can be connected through among-individual differences in behaviour at the earliest stages of life.

Animal personality has been related to habitat use in other taxa. More explorative juvenile lemon sharks (*Negaprion brevirostris*) for instance, took more risks than less explorative individuals by swimming further from the shores in a subpopulation with low predator abundance (Dhellemmes et al., 2021). This enabled them to forage more efficiently, at the cost of higher exposure to predators. Similarly, bold golden-mantel ground squirrels (*Callospermophilus lateralis*) had larger core areas and occupied more perches in their areas than their shy counterparts (Aliperti et al., 2021). Bonnot et al. (2015) found that roe deer (*Capreolus capreolus*) that reacted less actively during capture and handling, also tended to use open habitats more than conspecifics that reacted more actively at capture. However, these studies mainly focused on habitat use, whereas here we studied time investments regardless of habitat type or usage in a fairly homogenous environment. We show here that time spent scanning was repeatable, with 12% (in summer) and 17% (in autumn) of the variation in scanning time being attributable to the among-individual level. Time investments thus differed consistently among individuals, indicating that certain individuals, namely those that spend less time scanning, systematically have more time to allocate to foraging and subsequently, to gain more resources than other individuals.

The current scientific debate within the field of animal personality is focusing on whether among-individual differences in behaviour mostly reflect among-individual differences in resource allocation or acquisition (Laskowski et al., 2021; Haave-Audet et al., 2021). Recent meta-analyses seem to suggest the latter, since boldness is not associated with a survival cost overall (Moiron et al., 2020; Haave-Audet et al., 2021). These analyses, however, do not investigate the possible connection between resource allocation and acquisition. In this study, we show how individual fawns with a longer latency to leave at capture also allocate less time to scanning their environments months later during autumn, while in the herd with adult deer. Both behaviours could be classified as bold: individuals that stay during a capture conserve energy at the cost of risking mortality; likewise, individuals that allocate less time to scanning in the herd have more time to gain resources at the cost of predator detection. Thus, among-individual differences in behaviour can lead to differences in resource acquisition, through differences in allocation. Our results thereby provide a mechanistic pathway of among-individual differences that links allocation to acquisition.

The relationship between neonate capture response and time spent scanning was, contrary to our expectations, not present earlier on during summer. During this summer period, fawns make their first entrances into the herd with adult deer, after spending the first weeks of life hiding alone in the vegetation (Chapman & Chapman, 1997). In addition, fawns gradually switch from a nutrient rich diet (i.e. milk) to a nutrient poor diet (i.e. vegetation) with a concomitant need to invest more time in foraging. Fawns are thus very dependent on their mother for their resources during the first months and this dependency decreases with time, when their ability to forage successfully on their own becomes the main constraining factor for resource acquisition (Chapman & Chapman, 1997). As a result, scanning behaviour is expected to have a stronger limiting effect on foraging as fawns age, an effect clearly shown by our models. This suggests that scanning behaviour may not be functionally linked to life-history differences (here: resource acquisition) in summer, when fawns are also more dependent on milk of their mother, whereas this relationship is present in autumn. Therefore, even though the same behaviour was measured in summer and autumn, the functional role of that behaviour could be very different between life-stages. This may explain why we found no clear relationship between neonate personality and scanning behaviour in summer.

Another possibility is that relationships between different aspects of animal personality are overshadowed during major transitional phases in life. The emergence into the herd is such a major transition in the early life of fawns, where they are suddenly in the presence of many other conspecifics. From that point onwards, fawns socialise with other deer, and will therefore be exposed to many new stimuli. Dairy cattle, for instance, showed long-term consistency before and after puberty, but not across (Neave et al., 2020). Similarly, among-individual differences in red junglefowl (*Gallus gallus*) chicks’ behaviour were variable during ontogeny and stabilised after independence (Favati et al., 2016), a pattern also seen in wild fairy-wrens (*Malurus cyaneus*, Hall et al., 2015). On the other hand, there are also studies that do report long-term consistency across life-stages (Petelle et al., 2013; Debeffe et al., 2015). Petelle et al. (2013) show that yellow-bellied marmots (*Marmota flaviventris*) show long-term consistency in docility during captures, but not in boldness, whereas Debeffe et al. (2015) also show long-term consistency in docility, but then in wild roe deer (*Capreolus capreolus*) of which the youngest individuals were already months past their hiding phase. It is therefore possible that these studies found long-term consistency because they have not sampled individuals during transitional phases, but rather in between transitional phases.

Even though heart rates and latency to leave are strongly and inversely correlated in neonates at captures (Amin et al., 2021), we found no pattern between heart rates at capture and time spent scanning, suggesting that these two metrics are measuring separate traits. Captures of wild animals can be a stressful event, and typically evoke an acute stress response in prey animals such as fallow deer, which leads to an increased physiological and behavioural response (Harris & Carr, 2016). This relationship between physiology and behaviour does not have to be present at other times, such as during foraging bouts where animals are expected to have lower anxiety levels, and therefore also lower HPA-axis activation (Harris & Carr, 2016). Our findings in this study emphasise the need to include both physiological and behavioural responses to gain a better understanding of how physiology and behaviour are (or are not) related in different contexts.

Adult herbivores are classically expected to trade-off their time investments between anti-predator behaviour and resource acquisition. Although juveniles are not studied as extensively, previous research does indicate that juveniles differ from adults in the amount of time they spend scanning (Caro, 2005). In most birds and mammals, juveniles are shown to spend less time scanning than adults (e.g. Alados, 1985; Lashley et al., 2014; Li et al., 2015). The general explanation is that juveniles fail to recognize threats from predators and as a consequence spend less time scanning. In species where juveniles have a greater risk of being predated upon due to their reduced ability to escape, however, they may spend more time scanning due to the increased mortality threat (Caro, 2005). Our results show that fallow deer juveniles follow this pattern: as fawns grew older, they reduced their time spent scanning. This decrease in scanning time was accompanied by an increase in time spent foraging, a natural consequence of the weaning process (Chapman & Chapman, 1997).

To conclude, we have provided empirical support for the relationship between innate among-individual differences and resource acquisition, through allocation, suggesting a mechanistic pathway in which personality is associated with life-history. We have done so in juveniles of a wild large mammal, which have received little attention in the literature compared to other taxa (Bell et al., 2009). We furthermore have highlighted the development of among-individual variation from birth, throughout the transition from a solitary lifestyle to a group living one, up until the sixth month of life. Our results highlight how transitional phases can complicate patterns between behaviour and life-history, thereby offering novel insights into the ontogeny of animal personality. Overall, our study emphasizes the importance of including ontogeny for future studies, and the necessity to understand the relationship between allocation and acquisition for the improvement of theory in the field of animal personality.

## Acknowledgements

We thank the Office of Public Works (OPW), Ireland, for funding (grant no. R18625) and support. We extend a special thanks to Margaret Gormley (Chief Parks Superintendent), Paul McDonnell (Parks Superintendent), Maurice Cleary, Terry Moore, each of the OPW rangers, and the extended OPW staff in Phoenix Park – this study would not have been possible without them. We thank the School of Biology and Environmental Sciences (SBES) in University College Dublin (UCD) for co-funding this project. We like to thank Pippa-Jordan Faull, Sanne Fennema, May Higgins, Órla Heussaff, Ruairí O’Dea, Julia Evans, Kate Toland, Anthony Legeard and Sarah Keenen for their assistance in the field. The funders had no role in study design, data collection and analysis, decision to publish, or preparation of the manuscript. We declare that none of the authors have a conflict of interest.

## Data availability statement

Data and code (in R-Markdown format) will be published on Dryad upon acceptance and are already available as attachments to the editor and reviewers.

