## Supplementary material for "Neonate personality affects early-life resource acquisition in a large social mammal": S1

**Description of document:** This document contains details about the time that fawns emerge into the herd, after their hider-phase, and how this data was collected.

*Emergence into the herd*

Fallow deer fawns are introduced into the herd by their mother, at the end of their hiding phase, typically after at least a few weeks of life. The exact timing of emergence into the herd can be variable between individuals. Here we aimed at recording when each individual fawn entered the herd (i.e. herd emergence). Herd emergence was taken as the first day that an individual fawn was seen in a herd with an arbitrary size of at least 5 individuals, after being born (birthdays were derived from age estimates at capture, see Amin et al., 2021). For that purpose, we combined observations from four different data collections: Regular census surveys, summer herd presence, focal observations and presence data collected for a different study over the summer months (see Griffin et al., in prep). Data collection commenced on the 30$^{th}$ of July in 2018 and the 3$^{rd}$ of July in 2019.

As part of population monitoring, we have conducted regular census surveys from August 2018 onwards. These surveys took place at least once per month, with the exception of the month October, when the rutting season takes place. During these surveys, we walked through all the sectors in the park that are populated by the deer. We noted the GPS (brand) coordinates of each herd of deer encountered, including deer that were on their own. In addition, we noted the IDs of the individuals in these groups, the herd size and composition (sex and age). For each cohort, we took survey data up until the following fawning season in June.

During the period that we collected focal observation data (see below), we collected data on the location of individual fawns for both the focal observations and the summer presence. The summer presence data collection consisted of us recording the IDs, group size and location of every fawn we saw while surveying the park in search of fawns that could be sampled for the focal observations. Here we also recorded the IDs of fawns that were not sampled during the focal observations. The summer presence data was therefore done on the same days as the summer focal observations. Our final database consisted of 5626 observations (census survey: 2828; summer herd presence: 1184; focals: 898; parallel study, Griffin et al.: 716). This led to an herd emergence datum for 161 different fawns over two different years. These are shown in Figure S1.1

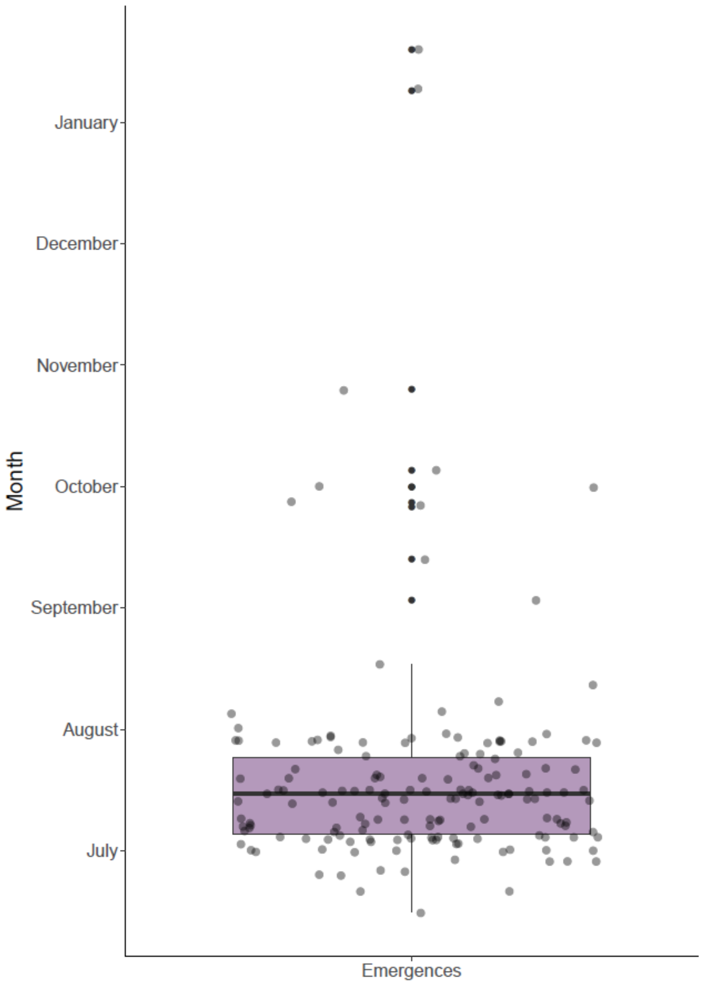


**Fig S1.1** Variation in time of emergence into the herd among 161 individual fawns over two subsequent years (2018 and 2019). Jittered points indicate individual fawns.
