## Supplementary material for "Neonate personality affects early-life resource acquisition in a large social mammal": ST1

**Description of document:** This document contains full details about all the behaviours recorded during focal time budget sampling, as reported in the main manuscript.

**Table S1:** Full list of behaviours recorded with description.

| Behaviour | Description |
|---|---|
| Grazing | Animal unselectively feeding on grass and ground vegetation, may be moving but activity is focused on ground with head below shoulder. |
| Browsing | Animal selectively feeding for leaves, bark and top of plants and shrubs. Head can be above the shoulders. |
| Scanning | Animal is standing straight up, head is erect above shoulder height. Eyes and ears focused, not chewing. |
| Scanning w/ chew | Animal is standing straight up, head erect and is chewing. |
| Suckling | Fawn is suckling milk from a doe, approached from the front and side. |
| Allosuckling | Fawn is suckling milk from a doe, approached from behind and positioned between the back legs of the doe |
| Self-Grooming | Animal is preening itself with muzzle. |
| Social Grooming (Giving) | Fawn is grooming another animal with muzzle. |
| Social Grooming (Receiving) | Fawn is being groomed by another animal, may still be carrying out other behaviours, such as grazing or scanning |
| Scratching | Fawn is scratching itself with own legs or off of other objects, such as trees. |
| Walking | Fawn is moving in slow pace, each leg moving in tangent. Fawn is not grazing or browsing. |
| Running | Fawn moving with increased speed, legs moving in tangent. |
| Pronking | Fawn moving quickly with all legs off the ground together. |
| Fighting | Fawn making contact with another animal, headbutting, pucking, pushing, shouldering, biting etc. |
| Object Play | Fawn manipulating and interacting with objects in the environment |
| Social Play | Fawn interacting with one or more other individuals |
| Locomotor Play | Fawn making rapid movements such as hops and springs. |
| Sniffing | Fawn is sniffing another deer |
| Flehman response | Fawn has head in the air and upper lip curled back |
| Digging | Fawn is digging up the ground with feet or with muzzle |
| Nudging | Fawn is nudging another animal with its head. The other animal may be lying or active, nudge may be light or forceful. |
| Vocalising Standing | Fawn is standing still and making noise |
| Vocalising Walking | Fawn is walking and making noise |
| Vocalising Running | The fawn is running and making noise |
| Mounting Giving | Fawn is mounting another deer |
| Mounting Receiving | Fawn is being mounted by another deer |
