## Supplementary material for "Neonate personality affects early-life resource acquisition in a large social mammal": S2: Supplementary_S2_Bivariate-models.html

S2: Full analysis of bivariate models reported in Amin et al.: Neonate personality affects early-life resource acquisition in a large social mammal


### S2: Full analysis of bivariate models reported in Amin et al.: Neonate personality affects early-life resource acquisition in a large social mammal

### Description:

This markdown contains the code to run the bivariate models as reported in the above paper, together with the full model outputs. The datasets that are used are created from the raw data (“Raw\_data.csv”), by use of the other script “SR1\_Create\_model\_datasets.R”.

This markdown is mainly divided into two sections: A) Creating final models and B) Results. Section A will use the datasets to create full models, then simplify them to the final models. Finally, diagnostics and model output will be checked. Section B will use the final models to reproduce the results reported in the main manuscript.

There is freedom to rerun models with the data and code provided here. In that case, we’d like to emphasize that the exact values of newly reran bivariate models may slightly differ from the values reported in the paper. This is entirely normal and due to the process of MCMC. However, all conclusions and main findings should be exaclty the same and should not be affected.

##### Description of variable names used in this script and which variables they represent in the paper:

meancap = Prior behaviour as described in the manuscript X..Deer = Number of deer in the group X..People = Number of people within 50 meters (excluding observers) Capture = Capture number, i.e. whether it was the first, second, third, fourth or fifth capture of the same animal BDAY = The birthday of the fawn, as numerical variable (1-365) Days = Day of the year, as numerical variable (1-365)

### Section A: Creating final models

  We start the process of creating our models by loading the package and the datasets needed for the analysis. We correct the variable structures where needed after which the data is ready. We then create our weakly informative prior, after which everything is ready for our bivariate models. We will go through each model one by one, starting with the full model and ending with the final model.

```
library(MCMCglmm)


Latency_Summer = read.csv("Latency_Summer_final.csv")
Latency_Autumn = read.csv("Latency_Autumn_final.csv")
HR_Summer = read.csv("HR_Summer_final.csv")
HR_Autumn = read.csv("HR_Autumn_final.csv")

Latency_Summer$Variable = as.factor(Latency_Summer$Variable)
Latency_Summer$Year = as.factor(Latency_Summer$Year)
Latency_Summer$Season = as.factor(Latency_Summer$Season)
Latency_Summer$Sex = as.factor(Latency_Summer$Sex)
Latency_Summer$Variable = as.factor(ifelse(Latency_Summer$Variable == "Vigilance", "Scanning", as.character(Latency_Summer$Variable)))

Latency_Autumn$Variable = as.factor(Latency_Autumn$Variable)
Latency_Autumn$Year = as.factor(Latency_Autumn$Year)
Latency_Autumn$Season = as.factor(Latency_Autumn$Season)
Latency_Autumn$Sex = as.factor(Latency_Autumn$Sex)
Latency_Autumn$Variable = as.factor(ifelse(Latency_Autumn$Variable == "Vigilance", "Scanning", as.character(Latency_Autumn$Variable)))

HR_Summer$Variable = as.factor(HR_Summer$Variable)
HR_Summer$Year = as.factor(HR_Summer$Year)
HR_Summer$Season = as.factor(HR_Summer$Season)
HR_Summer$Sex = as.factor(HR_Summer$Sex)
HR_Summer$Variable = as.factor(ifelse(HR_Summer$Variable == "Vigilance", "Scanning", as.character(HR_Summer$Variable)))

HR_Autumn$Variable = as.factor(HR_Autumn$Variable)
HR_Autumn$Year = as.factor(HR_Autumn$Year)
HR_Autumn$Season = as.factor(HR_Autumn$Season)
HR_Autumn$Sex = as.factor(HR_Autumn$Sex)
HR_Autumn$Variable = as.factor(ifelse(HR_Autumn$Variable == "Vigilance", "Scanning", as.character(HR_Autumn$Variable)))

# Setting (uninformative) prior :

prior = list(R = list(V = diag(2), nu = 0.002), G = list(G1 = list(V = diag(2), nu = 1.002)))
```

##### A.1) Heart rate-scanning summer model

We provide the code for the full model below.

```
HR_Summer_model<- MCMCglmm(Response~(Variable-1)+
                       at.level(Variable,1):scale(poly(meancap,2), scale = TRUE) +
                       at.level(Variable,1):scale(poly(Weight,2), scale = TRUE) +
                       at.level(Variable,1):scale(poly(Time,2), scale = TRUE) +
                       at.level(Variable,1):Sex +
                       at.level(Variable,1):scale(Air.temperature, scale = TRUE) +
                       at.level(Variable,2):Season +
                       at.level(Variable,2):scale(poly(Time,2), scale = TRUE) +
                       at.level(Variable,2):scale(poly(X..People,2), scale = TRUE) +
                       at.level(Variable,2):scale(poly(X..Deer, 2), scale = TRUE) + 
                       at.level(Variable,2):scale(poly(BDAY,2), scale = TRUE) +
                       at.level(Variable,2):scale(poly(Days_in_herd,2), scale = TRUE) +
                       at.level(Variable,2):scale(poly(Duration,2), scale = TRUE) +
                       at.level(Variable,2):Sex, 
                     random=~us(Variable):FawnID, rcov=~idh(Variable):units, family= "gaussian", prior=prior,nitt=1050000,thin=500,burnin=50000, data=HR_Summer,verbose=TRUE, pr = TRUE)
```

We have saved the model chain we used. We will load that model now and inspect the output.

```
HR_Summer_model <- readRDS ("HR_Summer_model.rds")
summary(HR_Summer_model)
```

```
## 
##  Iterations = 50001:1049501
##  Thinning interval  = 500
##  Sample size  = 2000 
## 
##  DIC: 1874.84 
## 
##  G-structure:  ~us(Variable):FawnID
## 
##                                          post.mean l-95% CI u-95% CI eff.samp
## VariableHRend:VariableHRend.FawnID         0.29265  0.14304  0.45075     2000
## VariableScanning:VariableHRend.FawnID     -0.02937 -0.10066  0.04143     2140
## VariableHRend:VariableScanning.FawnID     -0.02937 -0.10066  0.04143     2140
## VariableScanning:VariableScanning.FawnID   0.10810  0.05335  0.17529     2000
## 
##  R-structure:  ~idh(Variable):units
## 
##                        post.mean l-95% CI u-95% CI eff.samp
## VariableHRend.units       0.5100   0.3739   0.6536     1956
## VariableScanning.units    0.8173   0.7132   0.9260     2000
## 
##  Location effects: Response ~ (Variable - 1) + at.level(Variable, 1):scale(poly(meancap, 2), scale = TRUE) + at.level(Variable, 1):scale(poly(Weight, 2), scale = TRUE) + at.level(Variable, 1):scale(poly(Time, 2), scale = TRUE) + at.level(Variable, 1):Sex + at.level(Variable, 1):scale(Air.temperature, scale = TRUE) + at.level(Variable, 2):Season + at.level(Variable, 2):scale(poly(Time, 2), scale = TRUE) + at.level(Variable, 2):scale(poly(X..People, 2), scale = TRUE) + at.level(Variable, 2):scale(poly(X..Deer, 2), scale = TRUE) + 
##     at.level(Variable, 2):scale(poly(BDAY, 2), scale = TRUE) + at.level(Variable, 2):scale(poly(Days_in_herd, 2), scale = TRUE) + at.level(Variable, 2):scale(poly(Duration, 2), scale = TRUE) + at.level(Variable, 2):Sex 
## 
##                                                                   post.mean  l-95% CI  u-95% CI eff.samp  pMCMC    
## VariableHRend                                                     -0.163194 -0.354794  0.049442     1839  0.127    
## VariableScanning                                                   0.007373 -0.187467  0.232439     2000  0.943    
## at.level(Variable, 1):scale(poly(meancap, 2), scale = TRUE)1       0.086235  0.013233  0.159804     1952  0.026 *  
## at.level(Variable, 1):scale(poly(meancap, 2), scale = TRUE)2      -0.070362 -0.153325  0.007517     2000  0.096 .  
## at.level(Variable, 1):scale(poly(Weight, 2), scale = TRUE)1        0.218865  0.140738  0.296181     2000 <5e-04 ***
## at.level(Variable, 1):scale(poly(Weight, 2), scale = TRUE)2        0.033062 -0.037092  0.101688     2000  0.365    
## at.level(Variable, 1):scale(poly(Time, 2), scale = TRUE)1          0.127308 -0.009462  0.242559     2000  0.050 .  
## at.level(Variable, 1):scale(poly(Time, 2), scale = TRUE)2          0.001400 -0.103583  0.105962     2000  0.979    
## at.level(Variable, 1):Sexm                                         0.239204 -0.033797  0.505479     2000  0.082 .  
## at.level(Variable, 1):scale(Air.temperature, scale = TRUE)         0.079555  0.001007  0.152289     2000  0.038 *  
## at.level(Variable, 2):SeasonSummer2018                            -0.469209 -0.718938 -0.207579     2147  0.001 ***
## scale(poly(Time, 2), scale = TRUE)1:at.level(Variable, 2)         -0.170305 -0.261236 -0.057655     2000  0.001 ***
## scale(poly(Time, 2), scale = TRUE)2:at.level(Variable, 2)          0.047465 -0.058186  0.149670     1716  0.357    
## at.level(Variable, 2):scale(poly(X..People, 2), scale = TRUE)1     0.079690  0.001717  0.156790     2000  0.037 *  
## at.level(Variable, 2):scale(poly(X..People, 2), scale = TRUE)2     0.009497 -0.066695  0.085969     2204  0.829    
## at.level(Variable, 2):scale(poly(X..Deer, 2), scale = TRUE)1       0.091509  0.010080  0.167330     2000  0.020 *  
## at.level(Variable, 2):scale(poly(X..Deer, 2), scale = TRUE)2      -0.016664 -0.103122  0.065367     2000  0.694    
## at.level(Variable, 2):scale(poly(BDAY, 2), scale = TRUE)1          0.027412 -0.059422  0.115153     2149  0.536    
## at.level(Variable, 2):scale(poly(BDAY, 2), scale = TRUE)2         -0.015486 -0.104239  0.087776     2000  0.743    
## at.level(Variable, 2):scale(poly(Days_in_herd, 2), scale = TRUE)1 -0.339940 -0.563461 -0.119433     2000 <5e-04 ***
## at.level(Variable, 2):scale(poly(Days_in_herd, 2), scale = TRUE)2 -0.071754 -0.248169  0.072327     2009  0.373    
## at.level(Variable, 2):scale(poly(Duration, 2), scale = TRUE)1      0.158230  0.073451  0.251238     2000 <5e-04 ***
## at.level(Variable, 2):scale(poly(Duration, 2), scale = TRUE)2     -0.171178 -0.249360 -0.095300     2000 <5e-04 ***
## Sexm:at.level(Variable, 2)                                        -0.289393 -0.513983 -0.079653     2000  0.014 *  
## ---
## Signif. codes:  0 '***' 0.001 '**' 0.01 '*' 0.05 '.' 0.1 ' ' 1
```

Based on this full model, we now create our simplified final models. We do so by removing the quadratic effects which have a p > 0.1. Those are “Days in herd”, “BDAY”, “X..Deer”, “X..People” & “Time”. This model is now as following:

```
R_HR_Summer_model<- MCMCglmm(Response~(Variable-1)+
                             at.level(Variable,1):scale(poly(meancap,2), scale = TRUE) +
                             at.level(Variable,1):scale(poly(Weight,2), scale = TRUE) +
                             at.level(Variable,1):scale(poly(Time,2), scale = TRUE) +
                             at.level(Variable,1):Sex +
                             at.level(Variable,1):scale(Air.temperature, scale = TRUE) +
                             at.level(Variable,2):Season +
                             at.level(Variable,2):scale(Time, scale = TRUE) +
                             at.level(Variable,2):scale(X..People, scale = TRUE) +
                             at.level(Variable,2):scale(X..Deer, scale = TRUE) + 
                             at.level(Variable,2):scale(BDAY, scale = TRUE) +
                             at.level(Variable,2):scale(Days_in_herd, scale = TRUE) +
                             at.level(Variable,2):scale(poly(Duration,2), scale = TRUE) +
                             at.level(Variable,2):Sex, 
                           random=~us(Variable):FawnID, rcov=~idh(Variable):units, family= "gaussian", prior=prior,nitt=1050000,thin=500,burnin=50000, data=HR_Summer,verbose=TRUE, pr = TRUE)
```

We now load our model chain that we used for the main results in the manuscript, after which we give a model summary.

```
R_HR_Summer_model <- readRDS ("R_HR_Summer_model.rds")
summary(R_HR_Summer_model)
```

```
## 
##  Iterations = 50001:1049501
##  Thinning interval  = 500
##  Sample size  = 2000 
## 
##  DIC: 1866.963 
## 
##  G-structure:  ~us(Variable):FawnID
## 
##                                          post.mean l-95% CI u-95% CI eff.samp
## VariableHRend:VariableHRend.FawnID         0.28904  0.14075  0.45320     2000
## VariableScanning:VariableHRend.FawnID     -0.03047 -0.09969  0.03834     2000
## VariableHRend:VariableScanning.FawnID     -0.03047 -0.09969  0.03834     2000
## VariableScanning:VariableScanning.FawnID   0.10917  0.04781  0.16980     2000
## 
##  R-structure:  ~idh(Variable):units
## 
##                        post.mean l-95% CI u-95% CI eff.samp
## VariableHRend.units       0.5124   0.3754   0.6607     1819
## VariableScanning.units    0.8109   0.6979   0.9235     2000
## 
##  Location effects: Response ~ (Variable - 1) + at.level(Variable, 1):scale(poly(meancap, 2), scale = TRUE) + at.level(Variable, 1):scale(poly(Weight, 2), scale = TRUE) + at.level(Variable, 1):scale(poly(Time, 2), scale = TRUE) + at.level(Variable, 1):Sex + at.level(Variable, 1):scale(Air.temperature, scale = TRUE) + at.level(Variable, 2):Season + at.level(Variable, 2):scale(Time, scale = TRUE) + at.level(Variable, 2):scale(X..People, scale = TRUE) + at.level(Variable, 2):scale(X..Deer, scale = TRUE) + at.level(Variable, 
##     2):scale(BDAY, scale = TRUE) + at.level(Variable, 2):scale(Days_in_herd, scale = TRUE) + at.level(Variable, 2):scale(poly(Duration, 2), scale = TRUE) + at.level(Variable, 2):Sex 
## 
##                                                                post.mean   l-95% CI   u-95% CI eff.samp  pMCMC    
## VariableHRend                                                 -1.636e-01 -3.564e-01  6.102e-02     2000  0.141    
## VariableScanning                                               3.341e-02 -1.532e-01  2.163e-01     2587  0.724    
## at.level(Variable, 1):scale(poly(meancap, 2), scale = TRUE)1   8.731e-02  6.876e-03  1.633e-01     2000  0.027 *  
## at.level(Variable, 1):scale(poly(meancap, 2), scale = TRUE)2  -6.940e-02 -1.465e-01  1.241e-02     2000  0.097 .  
## at.level(Variable, 1):scale(poly(Weight, 2), scale = TRUE)1    2.196e-01  1.436e-01  3.013e-01     2000 <5e-04 ***
## at.level(Variable, 1):scale(poly(Weight, 2), scale = TRUE)2    3.291e-02 -3.935e-02  1.089e-01     2000  0.383    
## at.level(Variable, 1):scale(poly(Time, 2), scale = TRUE)1      1.265e-01 -1.812e-03  2.448e-01     2000  0.043 *  
## at.level(Variable, 1):scale(poly(Time, 2), scale = TRUE)2     -9.068e-05 -1.020e-01  1.045e-01     1959  0.983    
## at.level(Variable, 1):Sexm                                     2.380e-01 -1.122e-02  5.012e-01     1862  0.076 .  
## at.level(Variable, 1):scale(Air.temperature, scale = TRUE)     7.958e-02  3.867e-03  1.538e-01     2189  0.041 *  
## at.level(Variable, 2):SeasonSummer2018                        -4.208e-01 -6.529e-01 -1.763e-01     2760 <5e-04 ***
## at.level(Variable, 2):scale(Time, scale = TRUE)               -1.588e-01 -2.517e-01 -6.045e-02     2000  0.002 ** 
## at.level(Variable, 2):scale(X..People, scale = TRUE)           7.205e-02 -1.246e-03  1.497e-01     2000  0.069 .  
## at.level(Variable, 2):scale(X..Deer, scale = TRUE)             8.809e-02  3.037e-03  1.634e-01     2000  0.036 *  
## at.level(Variable, 2):scale(BDAY, scale = TRUE)                3.122e-02 -5.365e-02  1.179e-01     2000  0.480    
## at.level(Variable, 2):scale(Days_in_herd, scale = TRUE)       -2.671e-01 -3.993e-01 -1.249e-01     2000 <5e-04 ***
## at.level(Variable, 2):scale(poly(Duration, 2), scale = TRUE)1  1.592e-01  7.811e-02  2.476e-01     1834 <5e-04 ***
## at.level(Variable, 2):scale(poly(Duration, 2), scale = TRUE)2 -1.734e-01 -2.563e-01 -9.229e-02     1861 <5e-04 ***
## Sexm:at.level(Variable, 2)                                    -3.192e-01 -5.549e-01 -1.152e-01     2500  0.007 ** 
## ---
## Signif. codes:  0 '***' 0.001 '**' 0.01 '*' 0.05 '.' 0.1 ' ' 1
```

We then check for chain convergence. We have ran separate chains which we load here, but feel free to run new chains of your own. instead.

```
mean(summary(R_HR_Summer_model)$Gcovariances[,4])
```

```
## [1] 2000
```

```
mean(abs(autocorr.diag(R_HR_Summer_model$VCV[,1:4], lag=c(1))))
```

```
## [1] 0.007110329
```

```
R_HR_Summer_model2 <- readRDS ("R_HR_Summer_model2.rds")
R_HR_Summer_model3 <- readRDS ("R_HR_Summer_model3.rds")
R_HR_Summer_model4 <- readRDS ("R_HR_Summer_model4.rds")

diag = gelman.diag(mcmc.list(R_HR_Summer_model$Sol, R_HR_Summer_model2$Sol, R_HR_Summer_model3$Sol, R_HR_Summer_model4$Sol ))
diag$mpsrf
```

```
## [1] 1.057686
```

Everything seems okay. We have finalized this model.

##### A.2) Latency-scanning summer model

We provide the code for the full model below.

```
Lat_Summer<- MCMCglmm(Response~(Variable-1)+
                        at.level(Variable,1):scale(meancap, scale = TRUE) +
                        at.level(Variable,1):scale(poly(Weight,2), scale = TRUE) +
                        at.level(Variable,1):Year +
                        at.level(Variable,1):scale(Capture, scale = FALSE) +
                        at.level(Variable,2):Season +
                        at.level(Variable,2):scale(poly(Time,2), scale = TRUE) +
                        at.level(Variable,2):scale(poly(X..People,2), scale = TRUE) +
                        at.level(Variable,2):scale(poly(X..Deer, 2), scale = TRUE) + 
                        at.level(Variable,2):scale(poly(BDAY,2), scale = TRUE) +
                        at.level(Variable,2):scale(poly(Days_in_herd,2), scale = TRUE) +
                        at.level(Variable,2):scale(poly(Duration,2), scale = TRUE) +
                        at.level(Variable,2):Sex, 
                      random=~us(Variable):FawnID, rcov=~idh(Variable):units, family= "gaussian", prior=prior,nitt=1050000,thin=500,burnin=50000, data=Latency_Summer,verbose=TRUE, pr = TRUE)
```

We have saved the model chain we used. We will load that model now and inspect the output.

```
Lat_Summer <- readRDS ("Lat_Summer_model.rds")
summary(Lat_Summer)
```

```
## 
##  Iterations = 50001:1049501
##  Thinning interval  = 500
##  Sample size  = 2000 
## 
##  DIC: 1862.553 
## 
##  G-structure:  ~us(Variable):FawnID
## 
##                                          post.mean l-95% CI u-95% CI eff.samp
## VariableLatency:VariableLatency.FawnID    0.244634  0.11976  0.38936     1990
## VariableScanning:VariableLatency.FawnID  -0.004031 -0.06828  0.06163     2000
## VariableLatency:VariableScanning.FawnID  -0.004031 -0.06828  0.06163     2000
## VariableScanning:VariableScanning.FawnID  0.105480  0.05237  0.16549     2000
## 
##  R-structure:  ~idh(Variable):units
## 
##                        post.mean l-95% CI u-95% CI eff.samp
## VariableLatency.units     0.5045   0.3834   0.6417     2000
## VariableScanning.units    0.8195   0.7071   0.9296     1751
## 
##  Location effects: Response ~ (Variable - 1) + at.level(Variable, 1):scale(meancap, scale = TRUE) + at.level(Variable, 1):scale(poly(Weight, 2), scale = TRUE) + at.level(Variable, 1):Year + at.level(Variable, 1):scale(Capture, scale = FALSE) + at.level(Variable, 2):Season + at.level(Variable, 2):scale(poly(Time, 2), scale = TRUE) + at.level(Variable, 2):scale(poly(X..People, 2), scale = TRUE) + at.level(Variable, 2):scale(poly(X..Deer, 2), scale = TRUE) + at.level(Variable, 2):scale(poly(BDAY, 2), scale = TRUE) + at.level(Variable, 
##     2):scale(poly(Days_in_herd, 2), scale = TRUE) + at.level(Variable, 2):scale(poly(Duration, 2), scale = TRUE) + at.level(Variable, 2):Sex 
## 
##                                                                   post.mean  l-95% CI  u-95% CI eff.samp  pMCMC    
## VariableLatency                                                   -0.640693 -1.204770 -0.059208     2000  0.031 *  
## VariableScanning                                                   0.003315 -0.207447  0.211146     1786  0.985    
## at.level(Variable, 1):scale(meancap, scale = TRUE)                -0.128475 -0.205349 -0.053218     2157  0.003 ** 
## at.level(Variable, 1):scale(poly(Weight, 2), scale = TRUE)1       -0.142296 -0.222796 -0.059066     2000  0.003 ** 
## at.level(Variable, 1):scale(poly(Weight, 2), scale = TRUE)2        0.063701 -0.009815  0.129098     2000  0.092 .  
## at.level(Variable, 1):Year2019                                     0.100248 -0.162881  0.353419     2000  0.444    
## at.level(Variable, 1):scale(Capture, scale = FALSE)               -0.169971 -0.309696 -0.027856     2743  0.022 *  
## at.level(Variable, 2):SeasonSummer2018                            -0.471942 -0.718694 -0.229114     2000 <5e-04 ***
## at.level(Variable, 2):scale(poly(Time, 2), scale = TRUE)1         -0.174855 -0.278111 -0.077037     2000 <5e-04 ***
## at.level(Variable, 2):scale(poly(Time, 2), scale = TRUE)2          0.049021 -0.050707  0.152713     2000  0.366    
## at.level(Variable, 2):scale(poly(X..People, 2), scale = TRUE)1     0.081191  0.004146  0.161343     2000  0.046 *  
## at.level(Variable, 2):scale(poly(X..People, 2), scale = TRUE)2     0.010169 -0.058292  0.096340     2000  0.816    
## at.level(Variable, 2):scale(poly(X..Deer, 2), scale = TRUE)1       0.093604  0.016084  0.178809     2000  0.024 *  
## at.level(Variable, 2):scale(poly(X..Deer, 2), scale = TRUE)2      -0.015275 -0.094623  0.074250     2000  0.704    
## at.level(Variable, 2):scale(poly(BDAY, 2), scale = TRUE)1          0.021821 -0.063592  0.111701     2000  0.639    
## at.level(Variable, 2):scale(poly(BDAY, 2), scale = TRUE)2         -0.017978 -0.111979  0.080771     2000  0.678    
## at.level(Variable, 2):scale(poly(Days_in_herd, 2), scale = TRUE)1 -0.336564 -0.555631 -0.119217     1842  0.003 ** 
## at.level(Variable, 2):scale(poly(Days_in_herd, 2), scale = TRUE)2 -0.068354 -0.219869  0.097321     2278  0.399    
## at.level(Variable, 2):scale(poly(Duration, 2), scale = TRUE)1      0.159659  0.078755  0.243662     2000 <5e-04 ***
## at.level(Variable, 2):scale(poly(Duration, 2), scale = TRUE)2     -0.174026 -0.249473 -0.087978     2000 <5e-04 ***
## at.level(Variable, 2):Sexm                                        -0.279185 -0.494531 -0.044395     2000  0.016 *  
## ---
## Signif. codes:  0 '***' 0.001 '**' 0.01 '*' 0.05 '.' 0.1 ' ' 1
```

Based on this full model, we now create our simplified final models. We do so by removing the quadratic effects which have a p > 0.1. Those are “Days in herd”, “BDAY”, “X..Deer”, “X..People” & “Time”. This model is now as following:

```
R_Lat_Summer<- MCMCglmm(Response~(Variable-1)+
                        at.level(Variable,1):scale(meancap, scale = TRUE) +
                        at.level(Variable,1):scale(poly(Weight,2), scale = TRUE) +
                        at.level(Variable,1):Year +
                        at.level(Variable,1):scale(Capture, scale = FALSE) +
                        at.level(Variable,2):Season +
                        at.level(Variable,2):scale(Time, scale = TRUE) +
                        at.level(Variable,2):scale(X..People, scale = TRUE) +
                        at.level(Variable,2):scale(X..Deer, scale = TRUE) + 
                        at.level(Variable,2):scale(BDAY, scale = TRUE) +
                        at.level(Variable,2):scale(Days_in_herd, scale = TRUE) +
                        at.level(Variable,2):scale(poly(Duration,2), scale = TRUE) +
                        at.level(Variable,2):Sex, 
                      random=~us(Variable):FawnID, rcov=~idh(Variable):units, family= "gaussian", prior=prior,nitt=1050000,thin=500,burnin=50000, data=Latency_Summer,verbose=TRUE, pr = TRUE)
```

We now load our model chain that we used for the main results in the manuscript, after which we give a model summary.

```
R_Lat_Summer <- readRDS ("R_Lat_Summer_model.rds")
summary(R_Lat_Summer)
```

```
## 
##  Iterations = 50001:1049501
##  Thinning interval  = 500
##  Sample size  = 2000 
## 
##  DIC: 1854.796 
## 
##  G-structure:  ~us(Variable):FawnID
## 
##                                          post.mean l-95% CI u-95% CI eff.samp
## VariableLatency:VariableLatency.FawnID    0.243765  0.11227  0.37829     2000
## VariableScanning:VariableLatency.FawnID  -0.003659 -0.07272  0.06185     2000
## VariableLatency:VariableScanning.FawnID  -0.003659 -0.07272  0.06185     2000
## VariableScanning:VariableScanning.FawnID  0.107888  0.05351  0.16869     2000
## 
##  R-structure:  ~idh(Variable):units
## 
##                        post.mean l-95% CI u-95% CI eff.samp
## VariableLatency.units     0.5049   0.3795   0.6406     2466
## VariableScanning.units    0.8105   0.7059   0.9190     2000
## 
##  Location effects: Response ~ (Variable - 1) + at.level(Variable, 1):scale(meancap, scale = TRUE) + at.level(Variable, 1):scale(poly(Weight, 2), scale = TRUE) + at.level(Variable, 1):Year + at.level(Variable, 1):scale(Capture, scale = FALSE) + at.level(Variable, 2):Season + at.level(Variable, 2):scale(Time, scale = TRUE) + at.level(Variable, 2):scale(X..People, scale = TRUE) + at.level(Variable, 2):scale(X..Deer, scale = TRUE) + at.level(Variable, 2):scale(BDAY, scale = TRUE) + at.level(Variable, 2):scale(Days_in_herd, 
##     scale = TRUE) + at.level(Variable, 2):scale(poly(Duration, 2), scale = TRUE) + at.level(Variable, 2):Sex 
## 
##                                                               post.mean  l-95% CI  u-95% CI eff.samp  pMCMC    
## VariableLatency                                               -0.641662 -1.236809 -0.070466     1567  0.031 *  
## VariableScanning                                               0.029464 -0.141352  0.204863     2000  0.732    
## at.level(Variable, 1):scale(meancap, scale = TRUE)            -0.127867 -0.209776 -0.054888     1852  0.002 ** 
## at.level(Variable, 1):scale(poly(Weight, 2), scale = TRUE)1   -0.143536 -0.229553 -0.064646     2000 <5e-04 ***
## at.level(Variable, 1):scale(poly(Weight, 2), scale = TRUE)2    0.065008 -0.009149  0.135376     1624  0.085 .  
## at.level(Variable, 1):Year2019                                 0.098733 -0.178085  0.355750     2043  0.467    
## at.level(Variable, 1):scale(Capture, scale = FALSE)           -0.170315 -0.316689 -0.023898     1440  0.019 *  
## at.level(Variable, 2):SeasonSummer2018                        -0.410735 -0.637799 -0.178171     2000  0.001 ***
## at.level(Variable, 2):scale(Time, scale = TRUE)               -0.161659 -0.259339 -0.062616     2296 <5e-04 ***
## at.level(Variable, 2):scale(X..People, scale = TRUE)           0.071216 -0.007012  0.141111     2000  0.068 .  
## at.level(Variable, 2):scale(X..Deer, scale = TRUE)             0.089242  0.010017  0.166090     2000  0.022 *  
## at.level(Variable, 2):scale(BDAY, scale = TRUE)                0.025993 -0.060629  0.111653     2189  0.541    
## at.level(Variable, 2):scale(Days_in_herd, scale = TRUE)       -0.263545 -0.393362 -0.133821     2130 <5e-04 ***
## at.level(Variable, 2):scale(poly(Duration, 2), scale = TRUE)1  0.158088  0.075091  0.240625     2000  0.001 ***
## at.level(Variable, 2):scale(poly(Duration, 2), scale = TRUE)2 -0.174627 -0.252676 -0.095991     2000 <5e-04 ***
## at.level(Variable, 2):Sexm                                    -0.310197 -0.511686 -0.099316     2000  0.004 ** 
## ---
## Signif. codes:  0 '***' 0.001 '**' 0.01 '*' 0.05 '.' 0.1 ' ' 1
```

We then check for chain convergence. We have ran separate chains which we load here, but feel free to run new chains of your own. instead.

```
mean(summary(R_Lat_Summer)$Gcovariances[,4])
```

```
## [1] 2000
```

```
mean(abs(autocorr.diag(R_Lat_Summer$VCV[,1:4], lag=c(1))))
```

```
## [1] 0.00840204
```

```
R_Lat_Summer2 <- readRDS ("R_Lat_Summer_model2.rds")
R_Lat_Summer3 <- readRDS ("R_Lat_Summer_model3.rds")
R_Lat_Summer4 <- readRDS ("R_Lat_Summer_model4.rds")

diag = gelman.diag(mcmc.list(R_Lat_Summer$Sol, R_Lat_Summer2$Sol, R_Lat_Summer3$Sol, R_Lat_Summer4$Sol ))
diag$mpsrf
```

```
## [1] 1.056611
```

Everything seems okay. We have finalized this model.

##### A.3) Heart rate-scanning autumn model

We provide the code for the full model below.

```
HR_Autumn_model<- MCMCglmm(Response~(Variable-1)+
                       at.level(Variable,1):scale(poly(meancap,2), scale = TRUE) +
                       at.level(Variable,1):scale(poly(Weight,2), scale = TRUE) +
                       at.level(Variable,1):scale(poly(Time,2), scale = TRUE) +
                       at.level(Variable,1):Sex +
                       at.level(Variable,1):scale(Air.temperature, scale = TRUE) +
                       at.level(Variable,2):Season +
                       at.level(Variable,2):scale(poly(Time,2), scale = TRUE) +
                       at.level(Variable,2):scale(poly(X..People,2), scale = TRUE) +
                       at.level(Variable,2):scale(poly(X..Deer, 2), scale = TRUE) + 
                       at.level(Variable,2):scale(poly(BDAY,2), scale = TRUE) +
                       at.level(Variable,2):scale(poly(Days_in_herd,2), scale = TRUE) +
                       at.level(Variable,2):scale(poly(Duration,2), scale = TRUE) +
                       at.level(Variable,2):Sex, 
                     random=~us(Variable):FawnID, rcov=~idh(Variable):units, family= "gaussian", prior=prior,nitt=1050000,thin=500,burnin=50000, data=HR_Autumn,verbose=TRUE, pr = TRUE)
```

We have saved the model chain we used. We will load that model now and inspect the output.

```
HR_Autumn_model <- readRDS ("HR_Autumn_model.rds")
summary(HR_Autumn_model)
```

```
## 
##  Iterations = 50001:1049501
##  Thinning interval  = 500
##  Sample size  = 2000 
## 
##  DIC: 1809.235 
## 
##  G-structure:  ~us(Variable):FawnID
## 
##                                          post.mean l-95% CI u-95% CI eff.samp
## VariableHRend:VariableHRend.FawnID        0.278333  0.13994  0.44821     2000
## VariableScanning:VariableHRend.FawnID     0.002334 -0.08144  0.07937     2229
## VariableHRend:VariableScanning.FawnID     0.002334 -0.08144  0.07937     2229
## VariableScanning:VariableScanning.FawnID  0.160294  0.07693  0.24869     2000
## 
##  R-structure:  ~idh(Variable):units
## 
##                        post.mean l-95% CI u-95% CI eff.samp
## VariableHRend.units       0.5224   0.3928   0.6710     2000
## VariableScanning.units    0.8015   0.6806   0.9218     2000
## 
##  Location effects: Response ~ (Variable - 1) + at.level(Variable, 1):scale(poly(meancap, 2), scale = TRUE) + at.level(Variable, 1):scale(poly(Weight, 2), scale = TRUE) + at.level(Variable, 1):scale(poly(Time, 2), scale = TRUE) + at.level(Variable, 1):Sex + at.level(Variable, 1):scale(Air.temperature, scale = TRUE) + at.level(Variable, 2):Season + at.level(Variable, 2):scale(poly(Time, 2), scale = TRUE) + at.level(Variable, 2):scale(poly(X..People, 2), scale = TRUE) + at.level(Variable, 2):scale(poly(X..Deer, 2), scale = TRUE) + 
##     at.level(Variable, 2):scale(poly(BDAY, 2), scale = TRUE) + at.level(Variable, 2):scale(poly(Position, 2), scale = TRUE) + at.level(Variable, 2):scale(poly(Days_in_herd, 2), scale = TRUE) + at.level(Variable, 2):scale(poly(Duration, 2), scale = TRUE) + at.level(Variable, 2):Sex 
## 
##                                                                    post.mean   l-95% CI   u-95% CI eff.samp  pMCMC    
## VariableHRend                                                     -0.1418102 -0.3444866  0.0557312     2000  0.154    
## VariableScanning                                                  -0.0301809 -0.2804046  0.1922649     2000  0.798    
## at.level(Variable, 1):scale(poly(meancap, 2), scale = TRUE)1       0.0925190  0.0099126  0.1695323     1651  0.017 *  
## at.level(Variable, 1):scale(poly(meancap, 2), scale = TRUE)2      -0.0398756 -0.1216473  0.0467261     2153  0.361    
## at.level(Variable, 1):scale(poly(Weight, 2), scale = TRUE)1        0.1895987  0.1060764  0.2654688     2000 <5e-04 ***
## at.level(Variable, 1):scale(poly(Weight, 2), scale = TRUE)2        0.0317378 -0.0483607  0.1017657     2000  0.419    
## at.level(Variable, 1):scale(poly(Time, 2), scale = TRUE)1          0.1175303 -0.0059993  0.2371389     2000  0.066 .  
## at.level(Variable, 1):scale(poly(Time, 2), scale = TRUE)2          0.0161073 -0.1062829  0.1312734     2219  0.787    
## at.level(Variable, 1):Sexm                                         0.2249015 -0.0246853  0.4843732     2000  0.090 .  
## at.level(Variable, 1):scale(Air.temperature, scale = TRUE)         0.0780127  0.0022238  0.1524143     1095  0.046 *  
## at.level(Variable, 2):SeasonAutumnwinter2018                       0.2812062  0.0420707  0.5450936     2000  0.028 *  
## scale(poly(Time, 2), scale = TRUE)1:at.level(Variable, 2)         -0.1443299 -0.2509739 -0.0440281     2000  0.011 *  
## scale(poly(Time, 2), scale = TRUE)2:at.level(Variable, 2)          0.0531357 -0.0411995  0.1352213     2000  0.236    
## at.level(Variable, 2):scale(poly(X..People, 2), scale = TRUE)1     0.0400909 -0.0352687  0.1121815     1872  0.308    
## at.level(Variable, 2):scale(poly(X..People, 2), scale = TRUE)2    -0.0110500 -0.0964669  0.0585776     2000  0.791    
## at.level(Variable, 2):scale(poly(X..Deer, 2), scale = TRUE)1       0.0415163 -0.0448675  0.1218999     1834  0.329    
## at.level(Variable, 2):scale(poly(X..Deer, 2), scale = TRUE)2       0.0775760 -0.0002400  0.1682855     2000  0.066 .  
## at.level(Variable, 2):scale(poly(BDAY, 2), scale = TRUE)1         -0.0071345 -0.0879641  0.0906881     1836  0.860    
## at.level(Variable, 2):scale(poly(BDAY, 2), scale = TRUE)2         -0.0711295 -0.1693985  0.0262085     2000  0.153    
## at.level(Variable, 2):scale(poly(Position, 2), scale = TRUE)1      0.0398348 -0.0436752  0.1194865     2280  0.323    
## at.level(Variable, 2):scale(poly(Position, 2), scale = TRUE)2      0.0784864  0.0007781  0.1629199     2000  0.058 .  
## at.level(Variable, 2):scale(poly(Days_in_herd, 2), scale = TRUE)1 -0.2520577 -0.3777911 -0.1002669     2000 <5e-04 ***
## at.level(Variable, 2):scale(poly(Days_in_herd, 2), scale = TRUE)2  0.0039603 -0.1505206  0.1316215     2000  0.943    
## at.level(Variable, 2):scale(poly(Duration, 2), scale = TRUE)1      0.0438111 -0.0732815  0.1416366     2000  0.404    
## at.level(Variable, 2):scale(poly(Duration, 2), scale = TRUE)2     -0.0527979 -0.1454083  0.0505245     2000  0.279    
## Sexm:at.level(Variable, 2)                                        -0.0594749 -0.2815345  0.1776871     2000  0.621    
## ---
## Signif. codes:  0 '***' 0.001 '**' 0.01 '*' 0.05 '.' 0.1 ' ' 1
```

Based on this full model, we now create our simplified final models. We do so by removing the quadratic effects which have a p > 0.1. Those are “Duration”, “Days in herd”, “BDAY”, “X..People”, “Time”. This model is now as following:

```
R_HR_Autumn_model<- MCMCglmm(Response~(Variable-1)+
                             at.level(Variable,1):scale(poly(meancap,2), scale = TRUE) +
                             at.level(Variable,1):scale(poly(Weight,2), scale = TRUE) +
                             at.level(Variable,1):scale(poly(Time,2), scale = TRUE) +
                             at.level(Variable,1):Sex +
                             at.level(Variable,1):scale(Air.temperature, scale = TRUE) +
                             at.level(Variable,2):Season +
                             at.level(Variable,2):scale(Time, scale = TRUE) +
                             at.level(Variable,2):scale(X..People, scale = TRUE) +
                             at.level(Variable,2):scale(poly(X..Deer, 2), scale = TRUE) + 
                             at.level(Variable,2):scale(BDAY, scale = TRUE) +
                             at.level(Variable,2):scale(poly(Position,2), scale = TRUE) +
                             at.level(Variable,2):scale(Days_in_herd, scale = TRUE) +
                             at.level(Variable,2):scale(Duration, scale = TRUE) +
                             at.level(Variable,2):Sex, 
                           random=~us(Variable):FawnID, rcov=~idh(Variable):units, family= "gaussian", prior=prior,nitt=1050000,thin=500,burnin=50000, data=HR_Autumn,verbose=TRUE, pr = TRUE)
```

We now load our model chain that we used for the main results in the manuscript, after which we give a model summary.

```
R_HR_Autumn_model <- readRDS ("R_HR_Autumn_model.rds")
summary(R_HR_Autumn_model)
```

```
## 
##  Iterations = 50001:1049501
##  Thinning interval  = 500
##  Sample size  = 2000 
## 
##  DIC: 1805.296 
## 
##  G-structure:  ~us(Variable):FawnID
## 
##                                          post.mean l-95% CI u-95% CI eff.samp
## VariableHRend:VariableHRend.FawnID        0.278231  0.13562  0.44921     2000
## VariableScanning:VariableHRend.FawnID     0.002588 -0.07969  0.08822     2000
## VariableHRend:VariableScanning.FawnID     0.002588 -0.07969  0.08822     2000
## VariableScanning:VariableScanning.FawnID  0.161033  0.07341  0.24925     2000
## 
##  R-structure:  ~idh(Variable):units
## 
##                        post.mean l-95% CI u-95% CI eff.samp
## VariableHRend.units       0.5211   0.3834   0.6581     2000
## VariableScanning.units    0.8006   0.6753   0.9191     2000
## 
##  Location effects: Response ~ (Variable - 1) + at.level(Variable, 1):scale(poly(meancap, 2), scale = TRUE) + at.level(Variable, 1):scale(poly(Weight, 2), scale = TRUE) + at.level(Variable, 1):scale(poly(Time, 2), scale = TRUE) + at.level(Variable, 1):Sex + at.level(Variable, 1):scale(Air.temperature, scale = TRUE) + at.level(Variable, 2):Season + at.level(Variable, 2):scale(Time, scale = TRUE) + at.level(Variable, 2):scale(X..People, scale = TRUE) + at.level(Variable, 2):scale(poly(X..Deer, 2), scale = TRUE) + at.level(Variable, 
##     2):scale(BDAY, scale = TRUE) + at.level(Variable, 2):scale(poly(Position, 2), scale = TRUE) + at.level(Variable, 2):scale(Days_in_herd, scale = TRUE) + at.level(Variable, 2):scale(Duration, scale = TRUE) + at.level(Variable, 2):Sex 
## 
##                                                                post.mean   l-95% CI   u-95% CI eff.samp  pMCMC    
## VariableHRend                                                 -0.1454252 -0.3421394  0.0490201     2000  0.153    
## VariableScanning                                              -0.0363974 -0.2533348  0.1864772     2000  0.722    
## at.level(Variable, 1):scale(poly(meancap, 2), scale = TRUE)1   0.0931051  0.0209163  0.1792839     2000  0.018 *  
## at.level(Variable, 1):scale(poly(meancap, 2), scale = TRUE)2  -0.0410376 -0.1276969  0.0402949     2000  0.336    
## at.level(Variable, 1):scale(poly(Weight, 2), scale = TRUE)1    0.1881079  0.1122363  0.2734573     2000 <5e-04 ***
## at.level(Variable, 1):scale(poly(Weight, 2), scale = TRUE)2    0.0337223 -0.0490560  0.1057641     2000  0.405    
## at.level(Variable, 1):scale(poly(Time, 2), scale = TRUE)1      0.1168435 -0.0002902  0.2388702     2000  0.060 .  
## at.level(Variable, 1):scale(poly(Time, 2), scale = TRUE)2      0.0162343 -0.1145198  0.1292828     2000  0.790    
## at.level(Variable, 1):Sexm                                     0.2206471 -0.0241559  0.4849826     2136  0.095 .  
## at.level(Variable, 1):scale(Air.temperature, scale = TRUE)     0.0785917 -0.0006143  0.1547678     1836  0.045 *  
## at.level(Variable, 2):SeasonAutumnwinter2018                   0.3044684  0.0628830  0.5664566     2000  0.017 *  
## at.level(Variable, 2):scale(Time, scale = TRUE)               -0.1479454 -0.2518611 -0.0430867     2000  0.011 *  
## at.level(Variable, 2):scale(X..People, scale = TRUE)           0.0482077 -0.0307574  0.1191929     2000  0.193    
## at.level(Variable, 2):scale(poly(X..Deer, 2), scale = TRUE)1   0.0497260 -0.0328783  0.1292676     2000  0.246    
## at.level(Variable, 2):scale(poly(X..Deer, 2), scale = TRUE)2   0.0754508 -0.0029359  0.1593349     2000  0.066 .  
## at.level(Variable, 2):scale(BDAY, scale = TRUE)               -0.0041802 -0.0973345  0.0814344     2000  0.935    
## at.level(Variable, 2):scale(poly(Position, 2), scale = TRUE)1  0.0389762 -0.0374015  0.1211870     2000  0.335    
## at.level(Variable, 2):scale(poly(Position, 2), scale = TRUE)2  0.0891490  0.0034409  0.1690073     2000  0.039 *  
## at.level(Variable, 2):scale(Days_in_herd, scale = TRUE)       -0.2383501 -0.3694540 -0.1042241     2000 <5e-04 ***
## at.level(Variable, 2):scale(Duration, scale = TRUE)            0.0131079 -0.0810454  0.1099542     2000  0.766    
## Sexm:at.level(Variable, 2)                                    -0.0767319 -0.3062755  0.1544725     2000  0.517    
## ---
## Signif. codes:  0 '***' 0.001 '**' 0.01 '*' 0.05 '.' 0.1 ' ' 1
```

We then check for chain convergence. We have ran separate chains which we load here, but feel free to run new chains of your own. instead.

```
mean(summary(R_HR_Autumn_model)$Gcovariances[,4])
```

```
## [1] 2000
```

```
mean(abs(autocorr.diag(R_HR_Autumn_model$VCV[,1:4], lag=c(1))))
```

```
## [1] 0.008923824
```

```
R_HR_Autumn_model2 <- readRDS ("R_HR_Autumn_model2.rds")
R_HR_Autumn_model3 <- readRDS ("R_HR_Autumn_model3.rds")
R_HR_Autumn_model4 <- readRDS ("R_HR_Autumn_model4.rds")

diag = gelman.diag(mcmc.list(R_HR_Autumn_model$Sol, R_HR_Autumn_model2$Sol, R_HR_Autumn_model3$Sol, R_HR_Autumn_model4$Sol ))
diag$mpsrf
```

```
## [1] 1.052638
```

Everything seems okay. We have finalized this model.

##### A.4) Latency-scanning autumn model

We provide the code for the full model below.

```
Lat_Autumn<- MCMCglmm(Response~(Variable-1)+
                        at.level(Variable,1):scale(meancap, scale = TRUE) +
                        at.level(Variable,1):scale(poly(Weight,2), scale = TRUE) +
                        at.level(Variable,1):Year +
                        at.level(Variable,1):scale(Capture, scale = FALSE) +
                        at.level(Variable,2):Season +
                        at.level(Variable,2):scale(poly(Time,2), scale = TRUE) +
                        at.level(Variable,2):scale(poly(X..People,2), scale = TRUE) +
                        at.level(Variable,2):scale(poly(X..Deer, 2), scale = TRUE) + 
                        at.level(Variable,2):scale(poly(BDAY,2), scale = TRUE) +
                        at.level(Variable,2):scale(poly(Days_in_herd,2), scale = TRUE) +
                        at.level(Variable,2):scale(poly(Duration,2), scale = TRUE) +
                        at.level(Variable,2):Sex, 
                      random=~us(Variable):FawnID, rcov=~idh(Variable):units, family= "gaussian", prior=prior,nitt=1050000,thin=500,burnin=50000, data=Latency_Autumn,verbose=TRUE, pr = TRUE)
```

We have saved the model chain we used. We will load that model now and inspect the output.

```
Lat_Autumn <- readRDS ("Lat_Autumn_model.rds")
summary(Lat_Autumn)
```

```
## 
##  Iterations = 50001:1049501
##  Thinning interval  = 500
##  Sample size  = 2000 
## 
##  DIC: 1803.124 
## 
##  G-structure:  ~us(Variable):FawnID
## 
##                                          post.mean l-95% CI u-95% CI eff.samp
## VariableLatency:VariableLatency.FawnID     0.27184  0.11435 0.415221     2000
## VariableScanning:VariableLatency.FawnID   -0.07313 -0.16048 0.003271     2000
## VariableLatency:VariableScanning.FawnID   -0.07313 -0.16048 0.003271     2000
## VariableScanning:VariableScanning.FawnID   0.16932  0.08042 0.259533     2000
## 
##  R-structure:  ~idh(Variable):units
## 
##                        post.mean l-95% CI u-95% CI eff.samp
## VariableLatency.units     0.5415   0.4160   0.6801     2499
## VariableScanning.units    0.7976   0.6775   0.9162     2000
## 
##  Location effects: Response ~ (Variable - 1) + at.level(Variable, 1):scale(meancap, scale = TRUE) + at.level(Variable, 1):scale(poly(Weight, 2), scale = TRUE) + at.level(Variable, 1):Year + at.level(Variable, 1):scale(Capture, scale = FALSE) + at.level(Variable, 2):Season + at.level(Variable, 2):scale(poly(Time, 2), scale = TRUE) + at.level(Variable, 2):scale(poly(X..People, 2), scale = TRUE) + at.level(Variable, 2):scale(poly(X..Deer, 2), scale = TRUE) + at.level(Variable, 2):scale(poly(BDAY, 2), scale = TRUE) + at.level(Variable, 
##     2):scale(poly(Position, 2), scale = TRUE) + at.level(Variable, 2):scale(poly(Days_in_herd, 2), scale = TRUE) + at.level(Variable, 2):scale(poly(Duration, 2), scale = TRUE) + at.level(Variable, 2):Sex 
## 
##                                                                    post.mean   l-95% CI   u-95% CI eff.samp  pMCMC    
## VariableLatency                                                   -0.9726731 -1.5149115 -0.4031739     2000  0.001 ***
## VariableScanning                                                  -0.0334098 -0.2633332  0.2102777     2000  0.775    
## at.level(Variable, 1):scale(meancap, scale = TRUE)                -0.1321971 -0.2091576 -0.0503917     2000  0.002 ** 
## at.level(Variable, 1):scale(poly(Weight, 2), scale = TRUE)1       -0.0892834 -0.1734235  0.0013726     2138  0.051 .  
## at.level(Variable, 1):scale(poly(Weight, 2), scale = TRUE)2        0.0665626 -0.0093984  0.1341296     1858  0.071 .  
## at.level(Variable, 1):Year2019                                     0.1441266 -0.1127126  0.3919252     2000  0.292    
## at.level(Variable, 1):scale(Capture, scale = FALSE)               -0.2488440 -0.3924089 -0.1013584     2000  0.001 ***
## at.level(Variable, 2):SeasonAutumnwinter2018                       0.2761703  0.0219035  0.5257306     1854  0.026 *  
## at.level(Variable, 2):scale(poly(Time, 2), scale = TRUE)1         -0.1436601 -0.2528297 -0.0375210     2000  0.012 *  
## at.level(Variable, 2):scale(poly(Time, 2), scale = TRUE)2          0.0482735 -0.0368147  0.1321693     2000  0.259    
## at.level(Variable, 2):scale(poly(X..People, 2), scale = TRUE)1     0.0418387 -0.0335785  0.1151527     2000  0.285    
## at.level(Variable, 2):scale(poly(X..People, 2), scale = TRUE)2    -0.0139064 -0.0910813  0.0640478     2000  0.727    
## at.level(Variable, 2):scale(poly(X..Deer, 2), scale = TRUE)1       0.0446876 -0.0403516  0.1230802     2164  0.298    
## at.level(Variable, 2):scale(poly(X..Deer, 2), scale = TRUE)2       0.0738571 -0.0073270  0.1598684     2000  0.099 .  
## at.level(Variable, 2):scale(poly(BDAY, 2), scale = TRUE)1         -0.0129955 -0.1059652  0.0766192     2000  0.781    
## at.level(Variable, 2):scale(poly(BDAY, 2), scale = TRUE)2         -0.0662129 -0.1637677  0.0356644     2397  0.197    
## at.level(Variable, 2):scale(poly(Position, 2), scale = TRUE)1      0.0391176 -0.0386271  0.1158442     2000  0.310    
## at.level(Variable, 2):scale(poly(Position, 2), scale = TRUE)2      0.0806716  0.0002825  0.1637818     2821  0.054 .  
## at.level(Variable, 2):scale(poly(Days_in_herd, 2), scale = TRUE)1 -0.2476064 -0.3810461 -0.1074091     2180 <5e-04 ***
## at.level(Variable, 2):scale(poly(Days_in_herd, 2), scale = TRUE)2 -0.0017724 -0.1229500  0.1343806     2000  0.995    
## at.level(Variable, 2):scale(poly(Duration, 2), scale = TRUE)1      0.0465337 -0.0561321  0.1630544     2000  0.406    
## at.level(Variable, 2):scale(poly(Duration, 2), scale = TRUE)2     -0.0475881 -0.1429424  0.0505988     2000  0.348    
## at.level(Variable, 2):Sexm                                        -0.0602366 -0.3035002  0.1525333     2000  0.614    
## ---
## Signif. codes:  0 '***' 0.001 '**' 0.01 '*' 0.05 '.' 0.1 ' ' 1
```

Based on this full model, we now create our simplified final models. We do so by removing the quadratic effects which have a p > 0.1. Those are “Duration”, “Days in herd”, “BDAY”, “X..People”, “Time”. This model is now as following:

```
R_Lat_Autumn<- MCMCglmm(Response~(Variable-1)+
                        at.level(Variable,1):scale(meancap, scale = TRUE) +
                        at.level(Variable,1):scale(poly(Weight,2), scale = TRUE) +
                        at.level(Variable,1):Year +
                        at.level(Variable,1):scale(Capture, scale = FALSE) +
                        at.level(Variable,2):Season +
                        at.level(Variable,2):scale(Time, scale = TRUE) +
                        at.level(Variable,2):scale(X..People, scale = TRUE) +
                        at.level(Variable,2):scale(poly(X..Deer, 2), scale = TRUE) + 
                        at.level(Variable,2):scale(BDAY, scale = TRUE) +
                        at.level(Variable,2):scale(poly(Position,2), scale = TRUE) +
                        at.level(Variable,2):scale(Days_in_herd, scale = TRUE) +
                        at.level(Variable,2):scale(Duration, scale = TRUE) +
                        at.level(Variable,2):Sex, 
                      random=~us(Variable):FawnID, rcov=~idh(Variable):units, family= "gaussian", prior=prior,nitt=1050000,thin=500,burnin=50000, data=Latency_Autumn,verbose=TRUE, pr = TRUE)
```

We now load our model chain that we used for the main results in the manuscript, after which we give a model summary.

```
R_Lat_Autumn <- readRDS ("R_Lat_Autumn_model.rds")
summary(R_Lat_Autumn)
```

```
## 
##  Iterations = 50001:1049501
##  Thinning interval  = 500
##  Sample size  = 2000 
## 
##  DIC: 1797.967 
## 
##  G-structure:  ~us(Variable):FawnID
## 
##                                          post.mean l-95% CI u-95% CI eff.samp
## VariableLatency:VariableLatency.FawnID     0.27125  0.12375 0.419928     2239
## VariableScanning:VariableLatency.FawnID   -0.07627 -0.15863 0.006081     2465
## VariableLatency:VariableScanning.FawnID   -0.07627 -0.15863 0.006081     2465
## VariableScanning:VariableScanning.FawnID   0.16641  0.08285 0.252735     2000
## 
##  R-structure:  ~idh(Variable):units
## 
##                        post.mean l-95% CI u-95% CI eff.samp
## VariableLatency.units     0.5374   0.4013   0.6638     2000
## VariableScanning.units    0.7963   0.6797   0.9154     2000
## 
##  Location effects: Response ~ (Variable - 1) + at.level(Variable, 1):scale(meancap, scale = TRUE) + at.level(Variable, 1):scale(poly(Weight, 2), scale = TRUE) + at.level(Variable, 1):Year + at.level(Variable, 1):scale(Capture, scale = FALSE) + at.level(Variable, 2):Season + at.level(Variable, 2):scale(Time, scale = TRUE) + at.level(Variable, 2):scale(X..People, scale = TRUE) + at.level(Variable, 2):scale(poly(X..Deer, 2), scale = TRUE) + at.level(Variable, 2):scale(BDAY, scale = TRUE) + at.level(Variable, 2):scale(poly(Position, 
##     2), scale = TRUE) + at.level(Variable, 2):scale(Days_in_herd, scale = TRUE) + at.level(Variable, 2):scale(Duration, scale = TRUE) + at.level(Variable, 2):Sex 
## 
##                                                               post.mean  l-95% CI  u-95% CI eff.samp  pMCMC    
## VariableLatency                                               -0.969441 -1.554010 -0.449680     2000 <5e-04 ***
## VariableScanning                                              -0.032930 -0.244608  0.192679     1845  0.759    
## at.level(Variable, 1):scale(meancap, scale = TRUE)            -0.134694 -0.213846 -0.060044     2000 <5e-04 ***
## at.level(Variable, 1):scale(poly(Weight, 2), scale = TRUE)1   -0.087338 -0.177912 -0.003930     2000  0.050 *  
## at.level(Variable, 1):scale(poly(Weight, 2), scale = TRUE)2    0.065938 -0.007631  0.141662     1604  0.079 .  
## at.level(Variable, 1):Year2019                                 0.138657 -0.127380  0.399427     2000  0.298    
## at.level(Variable, 1):scale(Capture, scale = FALSE)           -0.248852 -0.391232 -0.106875     2000  0.001 ***
## at.level(Variable, 2):SeasonAutumnwinter2018                   0.295456  0.055543  0.549565     1826  0.024 *  
## at.level(Variable, 2):scale(Time, scale = TRUE)               -0.145608 -0.253281 -0.044264     2000  0.005 ** 
## at.level(Variable, 2):scale(X..People, scale = TRUE)           0.049964 -0.024374  0.122924     2000  0.197    
## at.level(Variable, 2):scale(poly(X..Deer, 2), scale = TRUE)1   0.053868 -0.026903  0.130206     2000  0.180    
## at.level(Variable, 2):scale(poly(X..Deer, 2), scale = TRUE)2   0.074229 -0.010222  0.153329     2000  0.081 .  
## at.level(Variable, 2):scale(BDAY, scale = TRUE)               -0.010276 -0.096756  0.077348     2000  0.842    
## at.level(Variable, 2):scale(poly(Position, 2), scale = TRUE)1  0.034903 -0.041317  0.112670     2000  0.411    
## at.level(Variable, 2):scale(poly(Position, 2), scale = TRUE)2  0.088097  0.005164  0.172193     2000  0.043 *  
## at.level(Variable, 2):scale(Days_in_herd, scale = TRUE)       -0.237638 -0.366449 -0.114740     2000 <5e-04 ***
## at.level(Variable, 2):scale(Duration, scale = TRUE)            0.022313 -0.079412  0.117907     2000  0.651    
## at.level(Variable, 2):Sexm                                    -0.083362 -0.312756  0.116464     2000  0.471    
## ---
## Signif. codes:  0 '***' 0.001 '**' 0.01 '*' 0.05 '.' 0.1 ' ' 1
```

We then check for chain convergence. We have ran separate chains which we load here, but feel free to run new chains of your own. instead.

```
mean(summary(R_Lat_Autumn)$Gcovariances[,4])
```

```
## [1] 2292.334
```

```
mean(abs(autocorr.diag(R_Lat_Autumn$VCV[,1:4], lag=c(1))))
```

```
## [1] 0.01615174
```

```
R_Lat_Autumn2 <- readRDS ("R_Lat_Autumn_model2.rds")
R_Lat_Autumn3 <- readRDS ("R_Lat_Autumn_model3.rds")
R_Lat_Autumn4 <- readRDS ("R_Lat_Autumn_model4.rds")

diag = gelman.diag(mcmc.list(R_Lat_Autumn$Sol, R_Lat_Autumn2$Sol, R_Lat_Autumn3$Sol, R_Lat_Autumn4$Sol ))
diag$mpsrf
```

```
## [1] 1.059498
```

Everything seems okay. We have finalized this model. We have now finalized the models.

### Section B: Results

##### Repeatability

We’ll start with the repeatability estimates of the neonate capture traits. For these, we will use the autumn models since they have a bigger sample size.

```
# Heart rate 
Rpt_HR <- R_HR_Autumn_model$VCV[,"VariableHRend:VariableHRend.FawnID"]/(
  R_HR_Autumn_model$VCV[,"VariableHRend:VariableHRend.FawnID"] +
    R_HR_Autumn_model$VCV[,"VariableHRend.units"])

plot(Rpt_HR)
```

```
mean(Rpt_HR)
```

```
## [1] 0.3453849
```

```
HPDinterval(Rpt_HR)
```

```
##         lower     upper
## var1 0.181436 0.5129434
## attr(,"Probability")
## [1] 0.95
```

```
# Latency to leave
Rpt_Lat <- R_Lat_Autumn$VCV[,"VariableLatency:VariableLatency.FawnID"]/(
  R_Lat_Autumn$VCV[,"VariableLatency:VariableLatency.FawnID"] +
    R_Lat_Autumn$VCV[,"VariableLatency.units"])

plot(Rpt_Lat)
```

```
mean(Rpt_Lat)
```

```
## [1] 0.3330523
```

```
HPDinterval(Rpt_Lat)
```

```
##         lower     upper
## var1 0.171318 0.4772475
## attr(,"Probability")
## [1] 0.95
```

We then continue with the repeatability of scanning. Since we have run two summer models and two autumn models, we’ll compute them from both, for transparency purposes. The estimates are near identical between the models within each season.

```
# Scanning in summer, from the heart rate model.
Rpt_Summer_Scanning <- R_HR_Summer_model$VCV[,"VariableScanning:VariableScanning.FawnID"]/(
  R_HR_Summer_model$VCV[,"VariableScanning:VariableScanning.FawnID"] +
    R_HR_Summer_model$VCV[,"VariableScanning.units"])

plot(Rpt_Summer_Scanning)
```

```
mean(Rpt_Summer_Scanning)
```

```
## [1] 0.1182554
```

```
HPDinterval(Rpt_Summer_Scanning)
```

```
##           lower    upper
## var1 0.06152734 0.184619
## attr(,"Probability")
## [1] 0.95
```

```
# And also the estimates from the latency summer model. 
Rpt_Summer_Scanning <- R_Lat_Summer$VCV[,"VariableScanning:VariableScanning.FawnID"]/(
  R_Lat_Summer$VCV[,"VariableScanning:VariableScanning.FawnID"] +
    R_Lat_Summer$VCV[,"VariableScanning.units"])

mean(Rpt_Summer_Scanning)
```

```
## [1] 0.1170484
```

```
HPDinterval(Rpt_Summer_Scanning)
```

```
##           lower    upper
## var1 0.06440102 0.181233
## attr(,"Probability")
## [1] 0.95
```

```
# Scanning in autumn, from the heart rate model.
Rpt_Autumn_Scanning <- R_HR_Autumn_model$VCV[,"VariableScanning:VariableScanning.FawnID"]/(
  R_HR_Autumn_model$VCV[,"VariableScanning:VariableScanning.FawnID"] +
    R_HR_Autumn_model$VCV[,"VariableScanning.units"])

plot(Rpt_Autumn_Scanning)
```

```
mean(Rpt_Autumn_Scanning)
```

```
## [1] 0.1667865
```

```
HPDinterval(Rpt_Autumn_Scanning)
```

```
##           lower     upper
## var1 0.08567439 0.2513043
## attr(,"Probability")
## [1] 0.95
```

```
# And also the estimates from the latency Autumn model. 
Rpt_Autumn_Scanning <- R_Lat_Autumn$VCV[,"VariableScanning:VariableScanning.FawnID"]/(
  R_Lat_Autumn$VCV[,"VariableScanning:VariableScanning.FawnID"] +
    R_Lat_Autumn$VCV[,"VariableScanning.units"])

mean(Rpt_Autumn_Scanning)
```

```
## [1] 0.1722518
```

```
HPDinterval(Rpt_Autumn_Scanning)
```

```
##           lower     upper
## var1 0.09557595 0.2586695
## attr(,"Probability")
## [1] 0.95
```

##### Covariation at the among-individual level

Finally, we will compute the covariations at the among-individual level. We’ll do that for all four models, which is the last step in this analysis.

```
# Covariation between heart rate and summer scanning 
Covar_HR_Summer_Scanning <- R_HR_Summer_model$VCV[,"VariableScanning:VariableHRend.FawnID"]/
  (sqrt(R_HR_Summer_model$VCV[,"VariableScanning:VariableScanning.FawnID"])*
     sqrt(R_HR_Summer_model$VCV[,"VariableHRend:VariableHRend.FawnID"]))

mean(Covar_HR_Summer_Scanning)
```

```
## [1] -0.1690418
```

```
HPDinterval(Covar_HR_Summer_Scanning)
```

```
##           lower     upper
## var1 -0.5256038 0.2073023
## attr(,"Probability")
## [1] 0.95
```

```
plot(Covar_HR_Summer_Scanning)
```

```
# Covariation between latency and summer scanning
Covar_Latency_Summer_Scanning <- R_Lat_Summer$VCV[,"VariableScanning:VariableLatency.FawnID"]/
  (sqrt(R_Lat_Summer$VCV[,"VariableScanning:VariableScanning.FawnID"])*
     sqrt(R_Lat_Summer$VCV[,"VariableLatency:VariableLatency.FawnID"]))

mean(Covar_Latency_Summer_Scanning)
```

```
## [1] -0.02368563
```

```
HPDinterval(Covar_Latency_Summer_Scanning)
```

```
##           lower     upper
## var1 -0.4343497 0.3536965
## attr(,"Probability")
## [1] 0.95
```

```
plot(Covar_Latency_Summer_Scanning)
```

```
# Covariation between heart rate and autumn scanning 
Covar_HR_Autumn_Scanning <- R_HR_Autumn_model$VCV[,"VariableScanning:VariableHRend.FawnID"]/
  (sqrt(R_HR_Autumn_model$VCV[,"VariableScanning:VariableScanning.FawnID"])*
     sqrt(R_HR_Autumn_model$VCV[,"VariableHRend:VariableHRend.FawnID"]))

mean(Covar_HR_Autumn_Scanning)
```

```
## [1] 0.01352408
```

```
HPDinterval(Covar_HR_Autumn_Scanning)
```

```
##           lower     upper
## var1 -0.3549883 0.4171766
## attr(,"Probability")
## [1] 0.95
```

```
plot(Covar_HR_Autumn_Scanning)
```

```
# Covariation between latency and autumn scanning
Covar_Latency_Autumn_Scanning <- R_Lat_Autumn$VCV[,"VariableScanning:VariableLatency.FawnID"]/
  (sqrt(R_Lat_Autumn$VCV[,"VariableScanning:VariableScanning.FawnID"])*
     sqrt(R_Lat_Autumn$VCV[,"VariableLatency:VariableLatency.FawnID"]))

mean(Covar_Latency_Autumn_Scanning)
```

```
## [1] -0.3591116
```

```
HPDinterval(Covar_Latency_Autumn_Scanning)
```

```
##           lower       upper
## var1 -0.6694338 -0.02799195
## attr(,"Probability")
## [1] 0.95
```

```
plot(Covar_Latency_Autumn_Scanning)
```
