## Supplementary material for "Neonate personality affects early-life resource acquisition in a large social mammal": S3: Supplementary_S3_Sensitivity-Analysis.html

S3: Sensitivity analysis reported in Amin et al.: Neonate personality affects early-life resource acquisition in a large social mammal


### S3: Sensitivity analysis reported in Amin et al.: Neonate personality affects early-life resource acquisition in a large social mammal

##### Explanation

Here, our aim is to perform a sensitivity analysis on the two main behaviours of interest: Scanning and Foraging. We will be dividing the datasets into two seasons (Summer and Autumn), as explained in the main manuscript.

For each season, we will run a GAMM with the behaviour (either Scanning or Foraging) as response and the observationlength, rounded off to minutes, as explanatory variable. From the output of the GAMM, we can see how robust the variables are and if they can be included in our analysis.

First, we’ll start by loading the datasets and the packages required. We then proceed to do a logit-transformation on the data, so that we can use a Gaussian error distribution later on. Since the logit of zero is infinite, we have added a small value to both the numerator and denominator, as suggested by Warton & Hui (2011). Finally, we scale the response variable. This process is the same as the one used for the main analysis of the manuscript and is explained in more details there.

Reference: Warton, D. I., & Hui, F. K. (2011). The arcsine is asinine: the analysis of proportions in ecology. Ecology, 92(1), 3-10.

```
library(mgcv)
library(segmented)


SummerScanning = read.csv("Data_SA_SummerScanning.csv")
AutumnScanning = read.csv("Data_SA_AutumnScanning.csv")
SummerForaging = read.csv("Data_SA_SummerForaging.csv")
AutumnForaging = read.csv("Data_SA_AutumnForaging.csv")

logitTransform <- function(p) { log( (p+0.002) / ((1-p)+0.002) ) }


SummerScanning$Response = logitTransform(SummerScanning$Response)
SummerScanning$Response = scale(SummerScanning$Response)
SummerScanning$FawnID = as.factor(SummerScanning$FawnID)

AutumnScanning$Response = logitTransform(AutumnScanning$Response)
AutumnScanning$Response = scale(AutumnScanning$Response)
AutumnScanning$FawnID = as.factor(AutumnScanning$FawnID)

SummerForaging$Response = logitTransform(SummerForaging$Response)
SummerForaging$Response = scale(SummerForaging$Response)
SummerForaging$FawnID = as.factor(SummerForaging$FawnID)

AutumnForaging$Response = logitTransform(AutumnForaging$Response)
AutumnForaging$Response = scale(AutumnForaging$Response)
AutumnForaging$FawnID = as.factor(AutumnForaging$FawnID)
```

##### Structure of data

In this sensitivity analysis, we use the focal observation data reported in the main manuscript. Scanning and foraging data are always used as a response, and observation length (in minutes) is used as explanatory variable. We have created bins for every minute, where we have included every observation shorter than that value. For example, data where Minutes = 1, have all the data points with observation lengths of up to 1 minute. Likewise, data where Minutes = 10 have all the observations binned into it with observation lengths up to 10 minutes. This is because we’re interested to see when data stabilize.

### Scanning data

##### Summer

We first start with Scanning behaviour during summer. We can do a raw plot of the proportion of time individuals spend scanning versus the observation length.

```
boxplot(SummerScanning$Response ~ SummerScanning$Minutes,main = "Scanning in summer", ylab = "Proportion of time spent scanning", xlab = "Observation length")
```

Now let’s run the GAMM. What we should look for is whether the 95% CI interval overlaps with the y-intercept (the horizontal line at y=0). We’ll include FawnID as random effect and the season (i.e. year) as a fixed effect into the model.

```
model <- gam(Response ~ s(Minutes) + Season + s(FawnID, bs = "re"), data = SummerScanning, family = gaussian)
plot.gam(model, select = 1)
abline(h = 0)
```

Conclusion:

We can see that there is some underestimation of scanning in observations that last less than 5 minutes, since the 95% CI does not overlap with zero there.

After these few starting minutes, scanning behaviour stabilizes and remains stable for all other lengths.

##### Autumn

We then do Scanning behaviour during autumn. We can do a raw plot of the proportion of time individuals spend scanning versus the observation length. If we do that, we note the following:

```
boxplot(AutumnScanning$Response ~ AutumnScanning$Minutes,main = "Scanning in Autumn", ylab = "Proportion of time spent scanning", xlab = "Observation length")
```

Now let’s run the GAMM. What we should look for is whether the 95% CI interval overlaps with the y-intercept (the horizontal line at y=0). We’ll include FawnID as random effect and the season (i.e. year) as a fixed effect into the model.

```
model <- gam(Response ~ s(Minutes)+ Season + s(FawnID, bs = "re"), data = AutumnScanning, family = gaussian)
plot.gam(model, select = 1)
abline(h = 0)
```

Conclusion:

We can see that the 95% CI of the GAMM always overlaps zero, even in the shortest observations. This remains so also during longer observations. We conclude, based on this analysis, that scanning behaviour in autumn is stable enough and barely affected by observation length.

### Foraging data

##### Summer

We first start with foraging behaviour during summer. We can do a raw plot of the proportion of time individuals spend foraging versus the observation length. If we do that, we note the following:

```
boxplot(SummerForaging$Response ~ SummerForaging$Minutes,main = "Foraging in summer", ylab = "Proportion of time spent foraging", xlab = "Observation length")
```

Now let’s run the GAMM. What we should look for is whether the 95% CI interval overlaps with the y-intercept (the horizontal line at y=0). We’ll include FawnID as random effect and the season (i.e. year) as a fixed effect into the model.

```
model <- gam(Response ~ s(Minutes)+ Season + s(FawnID, bs = "re"), data = SummerForaging, family = gaussian)
plot.gam(model, select = 1)
abline(h = 0)
```

Conclusion:

Similar to scanning behaviour during summer, we can see that foraging during summer is also underestimated in short observations. The point at which the 95% CI overlaps with 0 is, however, higher (roughly around 8 minutes) which indicates that longer observations are needed. In addition to the underestimation in shorter observations, we can also see that there is also overestimation in longer observations (between ~17 and ~24 minutes). This indicates that foraging behaviour doesn’t really stabilize with increasing observation durations, but rather remains unstable.

##### Autumn

We finally analyse foraging behaviour during autumn. We can do a raw plot of the proportion of time individuals spend scanning versus the observation length. If we do that, we note the following:

```
boxplot(AutumnForaging$Response ~ AutumnForaging$Minutes,main = "Foraging in Autumn", ylab = "Proportion of time spent foraging", xlab = "Observation length")
```

Now let’s run the GAMM. What we should look for is whether the 95% CI interval overlaps with the y-intercept (the horizontal line at y=0). We’ll include FawnID as random effect and the season (i.e. year) as a fixed effect into the model.

```
model <- gam(Response ~ s(Minutes) + Season + s(FawnID, bs = "re"), data = AutumnForaging, family = gaussian)
plot.gam(model, select = 1)
abline(h = 0)
```

Conclusion:

We notice immediately that, contrary to the scanning behaviour, the foraging behaviour seems to be more unstable in autumn than in summer. There is clear underestimation up until ~13 minute observations and also clear overestimation above ~17 minutes.

These patterns indicate that foraging behaviour during autumn is strongly affected by observation length, and does not stabilize at any time.

### Final overall conclusions

The goal of the analysis here was to determine whether scanning and foraging behaviour data are robust enough over different observation lengths. The sensitivity analysis performed here shows that scanning behaviour is relatively stable, with some underestimation below 5 minutes only during summer. During autumn, scanning behaviour is mostly unaffected by observation length and therefore a stable and robust measure.

The foraging data fared less well. In addition to a similar underestimation for the summer data as the scanning data mentioned above, there was also overestimation of the behaviour for longer observations, indicating an increased instability. Furthermore, contrary to scanning data, foraging data got only worse over time, with autumn data only being stable between ~13 and ~17 minute observations. Anything below that was underestimated, whereas anything above that suffered from overestimation.

Based on these results, we have decided to exclude foraging behaviour and to only include scanning behaviour for our among-individual comparisons. Although we are aware that we cannot fully correct the slight underestimation of scanning behaviour during summer, we have decided to include observation duration as a covariate in the bivariate models, to at least take into account some of the variation caused by observation length.
